# Large scale variation in the rate of *de novo* mutation, base composition, divergence and diversity in humans

**DOI:** 10.1101/110452

**Authors:** Thomas Smith, Peter Arndt, Adam Eyre-Walker

**Affiliations:** School of Life Sciences University of Sussex Brighton BN1 9QG United Kingdom; Max Planck Institute for Molecular Genetics Ihnestr. 63/73 14195 Berlin Germany

## Abstract

It has long been suspected that the rate of mutation varies across the human genome at a large scale based on the divergence between humans and other species. It is now possible to directly investigate this question using the large number of *de novo* mutations (DNMs) that have been discovered in humans through the sequencing of trios. We show that there is variation in the mutation rate at the 100KB, 1MB and 10MB scale that cannot be explained by variation at smaller scales, however the level of this variation is modest at large scales – at the 1MB scale we infer that ~90% of regions have a mutation rate within 50% of the mean. Different types of mutation show similar levels of variation and appear to vary in concert which suggests the pattern of mutation is relatively constant across the genome and hence unlikely to generate variation in GC-content. We confirm this using two different analyses. We find that genomic features explain less than 50% of the explainable variance in the rate of DNM. As expected the rate of divergence between species and the level of diversity within humans are correlated to the rate of DNM. However, the correlations are weaker than if all the variation in divergence was due to variation in the mutation rate. We provide evidence that this is due the effect of biased gene conversion on the probability that a mutation will become fixed. We find no evidence that linked selection affects the relationship between divergence and DNM density. In contrast to divergence, we find that most of the variation in diversity can be explained by variation in the mutation rate. Finally, we show that the correlation between divergence and DNM density declines as increasingly divergent species are considered.

**Author summary:** Using a dataset of 40,000 *de novo* mutations we show that there is large-scale variation in the mutation rate at the 100KB and 1MB scale. We show that different types of mutation vary in concert and in a manner that is not expected to generate variation in base composition; hence mutation bias is not responsible for the large-scale variation in base composition that is observed across human chromosomes. As expected large-scale variation in the rate of divergence between species and the variation within species across the genome, are correlated to the rate of mutation, but the correlation between divergence and the mutation rate is not as strong as they could be. We show that biased gene conversion is responsible for weakening the correlation. In contrast we find that most of the variation across the genome in diversity can be explained by variation in the mutation rate. Finally, we show that the correlation between the rate of mutation in humans and the divergence between humans and other species, weakens as the species become more divergent.

## Introduction

Until recently, the distribution of germ-line mutations across the genome was studied using patterns of nucleotide substitution between species in putatively neutral sequences (see [1] for review of this literature), since under neutrality the rate of substitution should be equal to the mutation rate. However, the sequencing of hundreds of individuals and their parents has led to the discovery of thousands of germ-line *de novo* mutations (DNMs) in humans [2-6]; it is therefore possible to begin analysing the pattern of DNMs directly rather than inferring their patterns from substitutions. Initial analyses have shown that the rate of germ-line DNM increases with paternal age [4], a result that was never-the-less inferred by Haldane some 70 years ago [7], varies across the genome [5] and is correlated to a number of factors, including the time of replication [3], the rate of recombination [3], GC content [5] and DNA hypersensitivity [5].

Previous analyses have demonstrated that there is large scale variation in the rate of DNM in both the germ-line [3, 5] and the somatic tissue [8–12]. Here we focus exclusively on germ-line mutations. We use a collection of over 40,000 germ-line DNMs to address a range of questions pertaining to the large-scale distribution of DNMs. First, we quantify how much variation there is, and investigate whether the variation in the mutation rate at a large-scale can be explained in terms of variation at smaller scales. We also investigate to what extent the variation is correlated between different types of mutation, and to what extent it is correlated to a range of genomic variables.

We use the data to investigate a long-standing question – what forces are responsible for the large-scale variation in GC content across the human genome, the so called “isochore” structure [13]. It has been suggested that the variation could be due to mutation bias [14–18], natural selection [13, 19, 20], biased gene conversion [21–24], or a combination of all three forces [25]. There is now convincing evidence that biased gene conversion plays a role in the generating at least some of the variation in GC-content [26–28]. However, this does not preclude a role for mutation bias or selection. With a dataset of DNMs we are able to test explicitly whether mutation bias causes variation in GC-content.

The rate of divergence between species is known to vary across the genome at a large scale [1]. As expected this appears to be in part due to variation in the rate of mutation [3]. However, the rate of mutation at the MB scale is not as strongly correlated to the rate of nucleotide substitution between species as it could be if all the variation in divergence between 1MB windows was due to variation in the mutation rate [3]. Instead, the rate of divergence appears to correlate to the rate of recombination as well. This might be due to one, or a combination, of several factors. First, recombination might affect the probability that a mutation becomes fixed by the process of biased gene conversion (BGC) (review by [26]). Second, recombination can affect the probability that a mutation will be fixed by natural selection; in regions of high recombination deleterious mutations are less likely to be fixed, whereas advantageous mutations are more likely. Third, low levels of recombination can increase the effects of genetic hitch-hiking and background selection, both of which can reduce the diversity in the human-chimp ancestor, and the time to coalescence and the divergence between species. There is evidence of this effect in the divergence of humans and chimpanzees, because the divergence between these two species is lower nearer exons and other functional elements [29]. And fourth, the correlation of divergence to both recombination and DNM density might simply be due to limitations in multiple regression; spurious associations can arise if multiple regression is performed on two correlated variables that are not known without error. For example, it might be that divergence only depends on the mutation rate, but that the mutation rate is partially dependent on the rate of recombination. In a multiple regression, divergence might come out as being correlated to both DNM density and the recombination rate, because we do not know the mutation rate without error, since we only have limited number of DNMs. Here, we introduce a test that can resolve between these explanations.

As with divergence, we might expect variation in the level of diversity across a genome to correlate to the mutation rate. The role of the mutation rate variation in determining the level of genetic diversity across the genome has long been a subject of debate. It was noted many years ago that diversity varies across the human genome at a large scale and that this variation is correlated to the rate of recombination [30–32]. Because the rate of substitution between species is also correlated to the rate of recombination, Hellmann et al. [30, 31] inferred that the correlation between diversity and recombination was at least in part due to a mutagenic effect of recombination. A number of recent studies have shown that recombination is mutagenic [3, 33, 34]. However, no investigation has recently been made as to whether this explains all the variation in diversity, or whether biased gene conversion, direct and linked selection have a major influence on diversity at a large scale.

## Results

### De novo mutations

To investigate large scale patterns of *de novo* mutations in humans we compiled data from four studies which between them had discovered 43,433 autosomal DNMs: 26,939 mutations from Wong et al. [6], 11016 mutations from Francioli et al. [3], 4931 mutations from Kong et al. [4] and 547 mutations from Michaelson et al. [5]. We divided the mutations up into 9 categories reflecting the fact that CpG dinucleotides have higher mutation rates than non-CpG sites, and the fact that we cannot differentiate which strand the mutation had occurred on: CpG C>T (a C to T or G to A mutation at a CpG site), CpG C>A, CpG C>G and for non-CpG sites C>T, T>C, C>A, T>G, C<>G and T<>A mutations.

The proportion of mutations in each category in each of the datasets is shown in figure 1. We find that the pattern of mutation differs significantly between the 4 studies (Chi-square test of independence on the number of mutations in each of the 9 categories, p < 0.0001). This appears to be largely due to the relative frequency of C>T transitions in both the CpG and non-CpG context. In the data from Wong et al. [6] and Michaelson et al. [5] the frequency of C>T transitions at CpG sites is ~13% whereas it is ~17% in the other two studies, a discrepancy which has been noted before [35, 36]. For non-CpG sites the frequency of C>T transitions is ~24% in all studies except that of Wong et al. in which it is 26%. It is not clear whether these patterns reflect differences in the mutation rate between different cohorts of individuals, possibly because of age [3, 4, 6] or geographical origin [37] or whether the differences are due to methodological problems associated with detecting DNMs.

**Figure 1.**
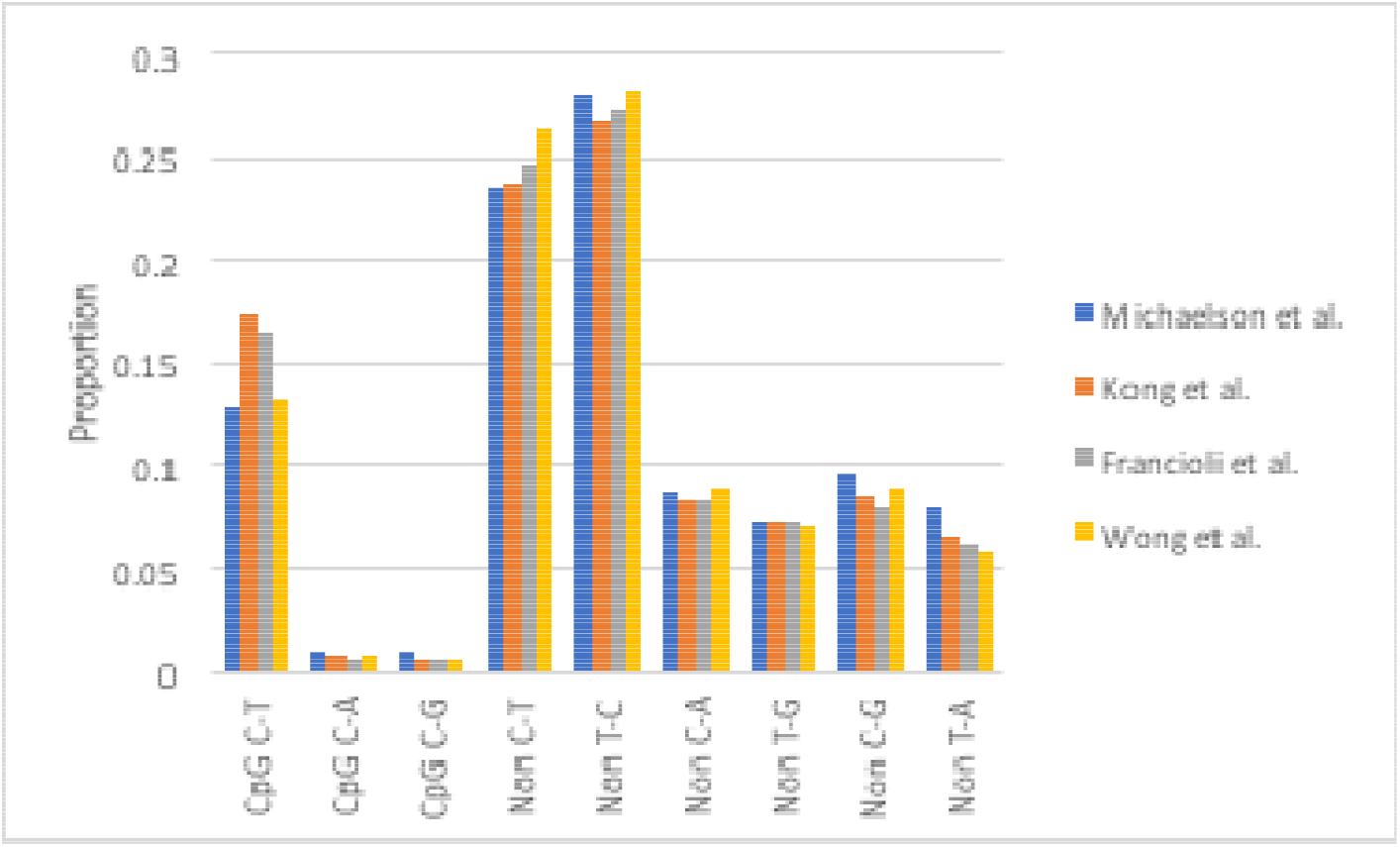
The proportion of DNMs in each of the mutational types in the four datasets.

### Distribution of rates

To investigate whether there is large scale variation in the mutation rate we divided the genome into non-overlapping windows of 10KB, 100KB, 1MB and 10MB and fit a gamma distribution to the number of mutations per region, taking into account the sampling error associated with the low number of mutations per region. We focussed our analysis at the 1MB scale since this has been extensively studied before. However, we show that the variation at 1MB forms part of a continuum of variation. We repeated almost all our analyses at the 100KB scale with qualitatively similar results (these results are reported in supplementary tables).

We find that the amount of variation differs significantly between the four studies (likelihood ratio tests: p < 0.001) with the level of variation being far greater in the Michaelson dataset, than in the Wong, Francioli and Kong datasets (Figure 2; Figure S1 for 100KB). The latter three datasets also show significantly different levels of variation (likelihood ratio test: p < 0.001). However, the differences between the three largest datasets are quantitatively small at the 1MB scale (Figure 2).

**Figure 2.**
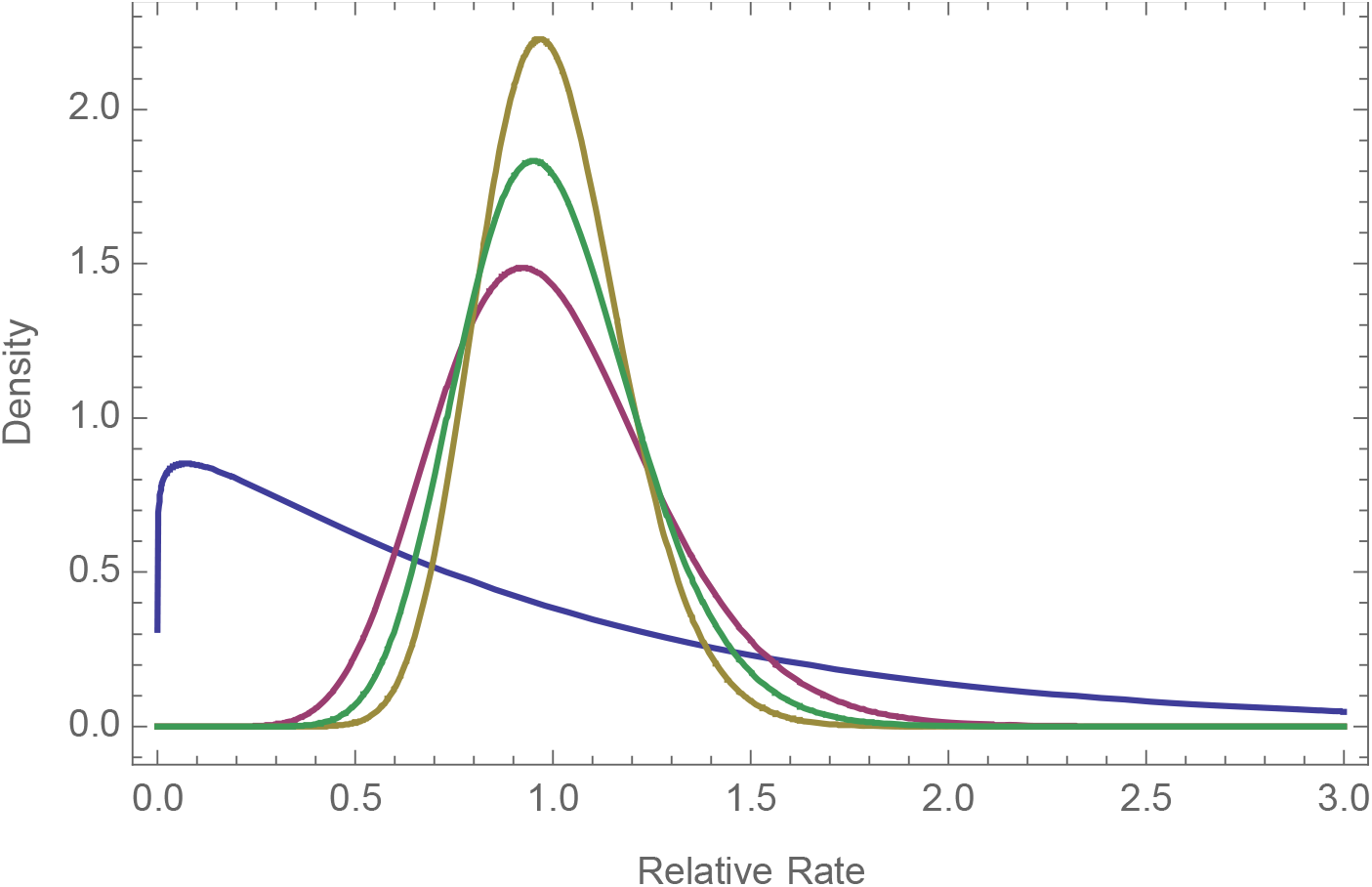
The gamma distribution fitted to the four datasets at the 1MB scales. Blue: Michaelson, Purple: Wong, Olive: Kong and Francioli (coincident distributions), Green: all data.

The variation between datasets might be due to differences in age or ethnicity between the individuals in each study, or methodological problems – for example, there might be differences between studies in the ability to identify DNMs. We can test whether callability is an issue in the largest of our datasets because Wong et al. [6] estimated the number of trios at which a DNM was callable at each site. If we reanalyse the Wong data using the sum of the callable trios per MB, rather than the number of sites in the human genome assembly, we obtain very similar estimates of the distribution: the coefficient of variation (CV) for the distribution is 0.27 when we use the number of sites and 0.25 when we use the sum of callable trios.

As expected the number of DNMs per site is significantly correlated between the Francioli and Wong datasets (1MB r =0.15, p<0.001). The correlation is weak, but this is likely to be in part due to sampling error. If we simulate data assuming a common distribution, estimating the shape parameter as the mean CV of the distributions fit to the individual datasets, the mean simulated correlation is 0.22. This suggests that much of the variation is common to the two datasets. However, subsequent analyses, reported below, show some important differences between the two datasets, and as a consequence we repeated all analyses on the Francioli and Wong datasets individually.

The CV of the gamma distribution fitted to the density of DNMs is 0.18 and 0.27 for the Francioli and Wong datasets respectively (Table 1). The level of variation is significant (i.e. the lower 95% confidence interval of the CV is greater than zero) (Table 1), however the level of variation is quite modest (Figure 3A, B). A gamma distribution with a coefficient of variation of 0.18 is one in which 90% of regions have a mutation rate within 30% of the mean (i.e. if the mean is one, between 0.7 and 1.3); for a gamma with a CV of 0.27, 95% of regions lie within 43% of the mean. The gamma distribution fits the distribution of rates qualitatively well (Figure S2), even though a goodness-of-fit test rejects the model at both the 100KB and 1MB scales (p<0.001). At both the 1MB and 100KB scales, there are some regions that have more DNMs than expected under the gamma distribution, and at the 1MB scale there are too many regions with no DNMs.

**Figure 3.**
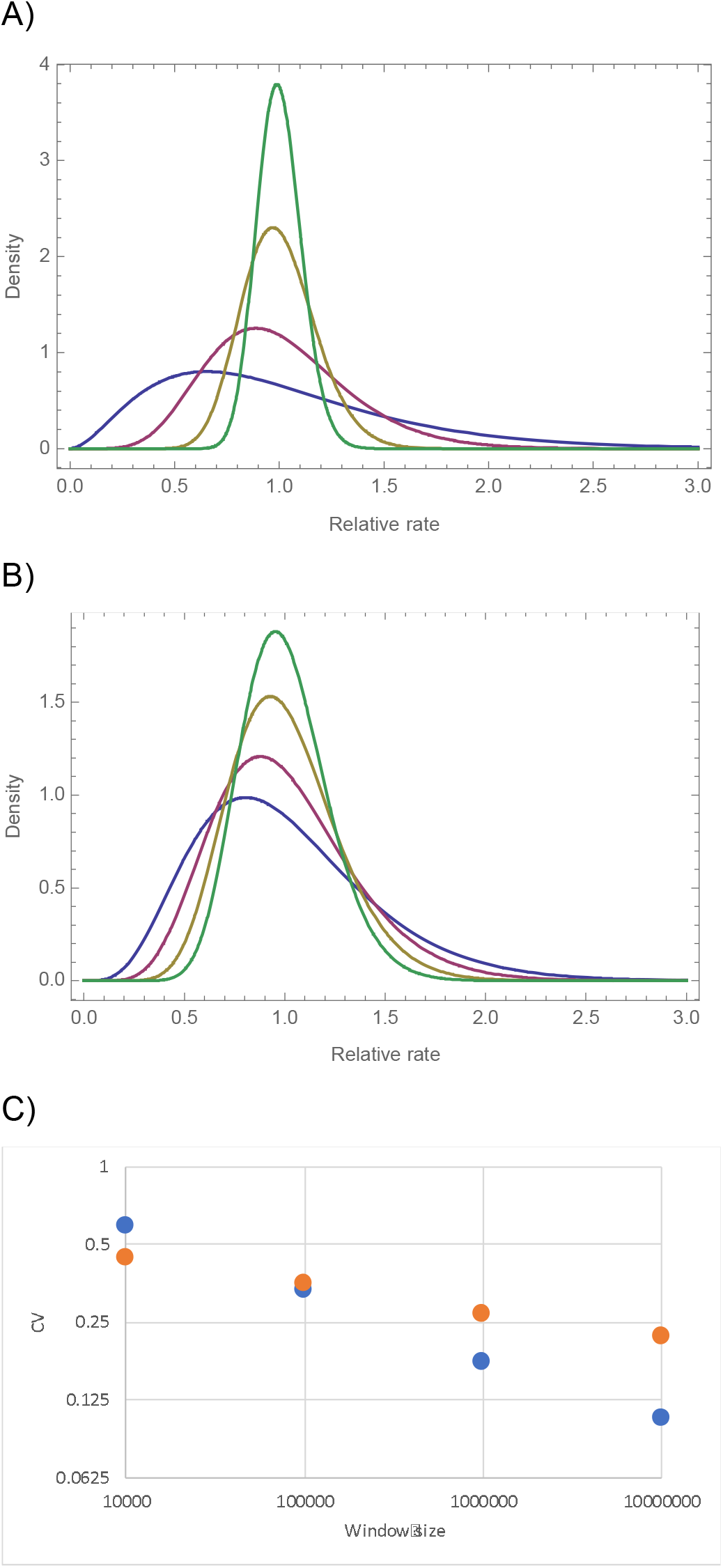
The gamma distribution fitted to the number of DNMs per window at the 10KB (blue) 100KB (purple), 1MB (olive) and 10MB (green) for the A) Francioli data and B) Wong data, and C) the coefficient of variation plotted against the window size, both on a log scale, for the Francioli (blue) and Wong (orange) datasets separately.

**Table 1.** The coefficient of variation for a gamma distribution fitted to the density of DNMs at the 1MB scale, and the 95% confidence intervals of the coefficient of variation. Strong(S) refers to G:C base pairs Weak(W) to A:T.

| Mutation type | Francioli | Wong |
| --- | --- | --- |
| Overall | 0.18 (0.13, 0.22) | 0.27 (0.25, 0.29) |
| CpG | 0.46 (0.35, 0.55) | 0.29 (0.21, 0.36) |
| nonCpG | 0.17 (0.11, 0.22) | 0.26 (0.24, 0.28) |
| CpG transitions | 0.43 (0.30, 0.53) | 0.29 (0.20, 0.36) |
| CpG transversions | 0.50 (0, 0.56) | 0.53 (0, 0.88) |
| nonCpG transitions | 0.20 (0.10, 0.26) | 0.24 (0.20, 0.26) |
| nonCpG transversions | 0.20 (0, 0.30) | 0.29 (0.25, 0.33) |
| S>W | 0.25 (0.19, 0.32) | 0.25 (0.22, 0.28) |
| W>S | 0.063 (0, 0.21) | 0.24 (0.20, 0.28) |

If we include estimates of the distribution for 10KB, 100KB and 10MB we find that the log of the CV of the gamma distribution is approximately linearly related to the log of the scale (Figure 3C). This suggests that the variation at the 1MB scale is part of a continuum of variation at different scales. The linearity of the relationship suggests that a simple phenomenon underlies the variation. However, the slope of this relationship differs between the Francioli (slope = -0.25) and Wong (-0.10) datasets suggesting that there are some systematic differences between these two studies.

If all the variation at the larger scales is explainable by variation at a smaller scale, then the CV at scale *x* should be equal to the CV at some finer scale, *y*, divided by the square-root of *x/y*; on a log-log scale this should yield a slope of -0.5. This therefore suggests that there is variation at a larger scale that cannot be explained by variation at smaller scale. To test whether this is the case, we ran a series of one-way ANOVAs; in all comparisons, testing variation at the 100KB scale using 10KB windows, 1MB using 100KB windows and 10MB using 1MB windows the results were significant for the combined data and for the Wong and Francioli datasets individually (p<0.001 in all cases).

### Mutational types

If we estimate the distribution for individual mutational types we find that in many cases the lower CI on the CV is zero; this might be because we do not have enough data to reliably estimate the distribution for each individual mutational type. We therefore combined mutations into a variety of nonmutually exclusive categories. In each case we estimated the distribution for the relevant category of sites – e.g. in considering the distribution of CpG rates we consider the number of CpG DNMs at CpG sites, not at all sites. We find that the estimated distributions are similar for different mutational types although there is rather more variation at CpG sites in the Francioli dataset (Table 1; 100KB results Table S1). Although the distributions are fairly similar for different mutational types, likelihood ratio tests demonstrate that there is significantly more variation at CpG sites than nonCpG sites and for S>W than W>S changes in the Francioli dataset (Table S2 for 1MB and 100KB results). We also find significant differences between non-CpG transitions and transversions at 100KB scale in both datasets. Never-the-less the differences between different mutational categories are relatively small.

### Correlations between mutational types

Given that there is variation in the mutation rate at the 1MB scale and that this variation is quite similar in magnitude for different mutational types, it would seem likely that the rate of mutation for the different mutational types are correlated. We find that this is indeed the case. We observe significant correlations between all categories of mutations in both the Francioli and Wong datasets, except between CpG transitions and transversions in the Wong dataset (Table 2; Table S3 for 100KB). The correlations are weak but this is to be expected given the large level of sampling error. To compare the correlation to what we might expect if the two categories of mutation shared a common distribution and were perfectly correlated, we simulated data under a common distribution, estimating the CV of the common distribution as the mean of the distributions fitted to the two mutational categories. We find that generally the observed correlations are similar, and not significantly different, to the expected correlations.

**Table 2.** The correlation between different mutational types at the 1MB scales. The observed correlation is given along with the mean correlation from data simulated under the assumption that the two categories have the same distribution and are perfectly correlated. * 0<0.05, **p<0.01, ***p<0.001

| Comparison | Observed correlation | Expected correlation | Proportion of simulated correlations > observed |
| --- | --- | --- | --- |
| *Francioli* |  |  |  |
| CpG v. nonCpG | 0.097*** | 0.11 | 0.72 |
| CpG ts v. CpG tv | 0.039* | 0.028 | 0.3 |
| nonCpG ts v. nonCpG tv | 0.052** | 0.062 | 0.57 |
| S>W v. W>S | 0.061** | 0.037 | 0.10 |
| *Wong* |  |  |  |
| CpG v. nonCpG | 0.16*** | 0.17 | 0.65 |
| CpG ts v. CpG tv | 0.035 | 0.053 | 0.77 |
| nonCpG ts v. nonCpG tv | 0.22*** | 0.20 | 0.23 |
| S>W v. W>S | 0.22*** | 0.19 | 0.15 |

### Variation in base composition

The fact that the rates of S>W and W>S mutation covary suggests that mutational biases are unlikely to generate much variation in GC-content across the genome. To investigate this further, we used two approaches to test whether there was variation in the pattern of mutation that could generate variation in GC content. First, we used the DNM data for each window to predict the equilibrium GC content to which the sequence would evolve, fitting a model by maximum likelihood (ML) in which this equilibrium GC-content could vary across the genome. For the Francioli data the ML model is one in which the equilibrium GC content has a mean of 0.32 and a standard deviation of 0.02. The Wong dataset yields similar estimates but the ML estimate of the standard deviation is 0.001. In both cases, the upper estimate on the standard deviation is small – 0.060 in Francioli and 0.036 in Wong suggesting that variation in the pattern of mutation is unlikely to generate much variation in GC content.

However, the ML method does not rule out the possibility that there is some variation in the pattern of mutation. Furthermore, the method does not take into account the difference in the mutation rate between CpG and non-CpG sites. We therefore used a second approach in which we ranked regions by their current GC-content and grouped regions together. We then estimated the mutation rates for all categories of mutation using the DNM data and used these estimated mutation rates in a simulation of sequence evolution, in which we evolved the sequence to its equilibrium GC content. We find no correlation between the equilibrium GC content to which the sequence evolves and the current GC content (Figure 4; Figure S3 for 100KB).

**Figure 4.**
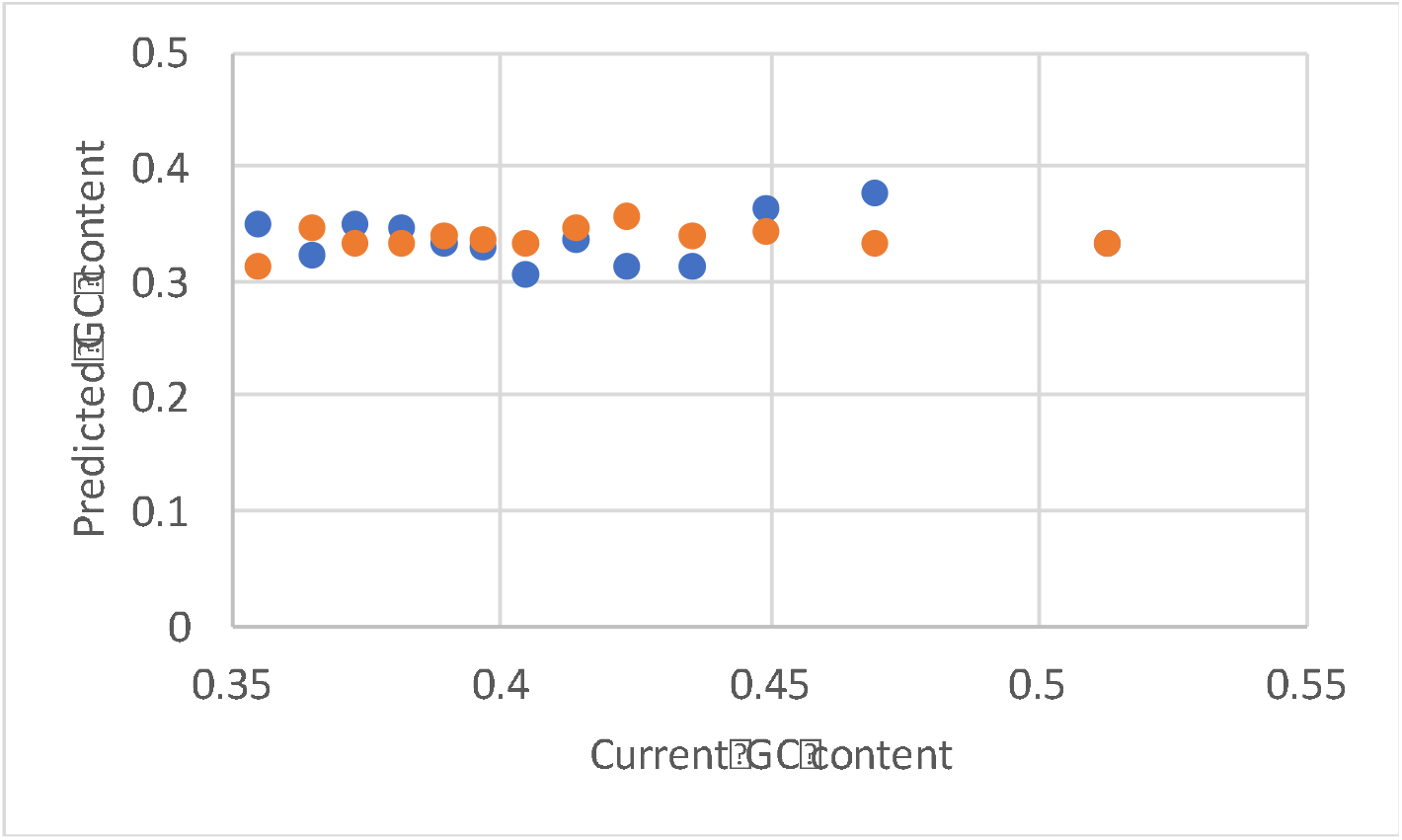
The predicted equilibrium GC content against the current GC content for the Francioli (blue) and Wong (orange) datasets. Note the right most point is coincident for the two datasets.

### Mutation models

It has been suggested that the mutation rate at a site is predictable based on genomic features, such as replication time [5], or the 7-mer sequence in which a site is found [38]. To investigate whether these models can explain the variation at large scales we used the models to predict the average mutation rate for each 100KB or 1MB region and correlated these predictions against the observed number of DNMs per site.

We find that the density of DNMs is significantly correlated to the rates predicted under the 7-mer model of Aggarwala et al. [38]. This correlation is significantly positive for the combined and Wong datasets, as we might expect, but significantly negative for the Francioli dataset (Table 3; Table S4 for 100KB results). To compare these correlations to what we might expect if the Aggarwala model explained all the variation at large scales, we simulated the appropriate number of DNMs across the genome according to this model. The observed correlation is significantly smaller than the expected correlation for all datasets, however, the observed and expected correlations are quite similar for the Wong dataset suggesting that much of the variation in DNM density in this dataset is explainable by the model of Aggarwala et al. [38]

**Table 3.** Correlation between the density of DNMs and the mutation rate estimates of Aggarwala et al and Michaelson et al. at the 1MB scale. The expected values are the mean correlations observed from 1000 simulations *p < 0.05 **p < 0.01 *** p < 0.001

|  | Aggarwala |  | Michaelson |  |
| --- | --- | --- | --- | --- |
|  | Observed | Expected | Observed | Expected |
| Francioli | -0.16*** | 0.16*** | 0.084*** | 0.41*** |
| Wong | 0.18*** | 0.25*** | 0.018 | 0.58*** |

In contrast, although the density of DNMs is significantly positively correlated to the predictions of the Michaelson model for the Francioli dataset, but not for the Wong dataset, the correlation is substantially and significantly smaller than it could be if the model explained all the variation (Table 3; Table S4 for 100KB results).

### Correlations with genomic variables

To try and understand why there is large scale variation in the mutation rate, we compiled a number of genomic variables which have previously been shown to correlate to the rate of germline or somatic DNM, or divergence between species: male and female recombination rate, GC content, replication time, nucleosome occupancy, transcription level, DNA hypersensitivity and several histone methylation and acetylation marks [3, 5, 9, 39, 40].

Unfortunately, the Wong and Francioli datasets yield different patterns of correlation. The overall density of DNMs is significantly positively correlated to male and female recombination rates, but many of the other significant correlations are in opposite directions (Table 4; 100KB results Table S5).

**Table 4.**
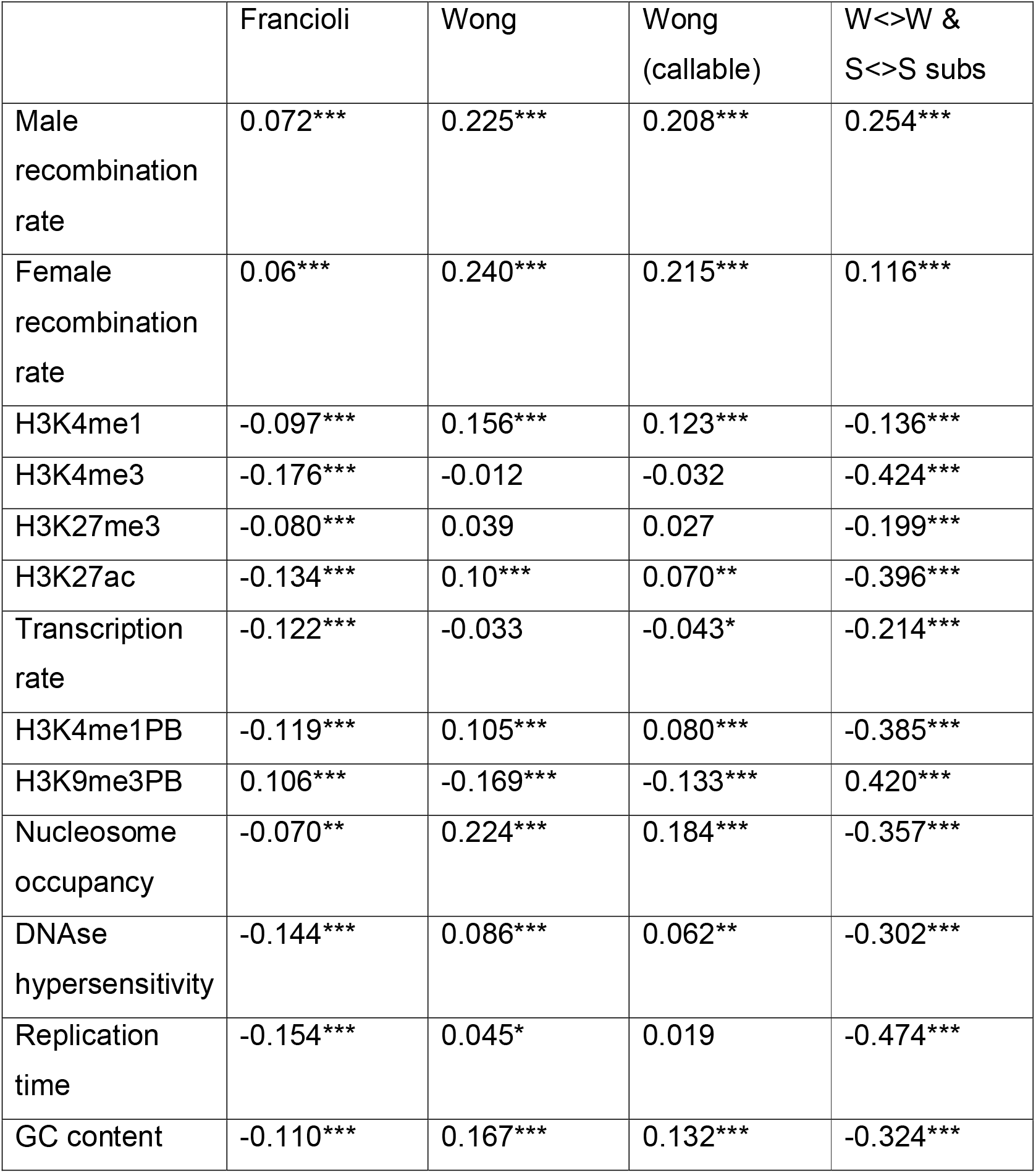
The correlation between the density of DNMs in the two largest datasets and various genomic variables at the 1MB scale. Also shown are the correlations when the number of DNMs in the Wong dataset divided by the sum of the callable trios and the number of W<>W and S<>S substitutions per site between human and chimpanzee. * p<0.05 **p<0.01 ***p<0.001.

|  | Francioli | Wong | Wong<br>(callable) | W<>W &<br>S<>S subs |
| --- | --- | --- | --- | --- |
| Male recombination rate | 0.072*** | 0.225*** | 0.208*** | 0.254*** |
| Female recombination rate | 0.06*** | 0.240*** | 0.215*** | 0.116*** |
| H3K4me1 | -0.097*** | 0.156*** | 0.123*** | -0.136*** |
| H3K4me3 | -0.176*** | -0.012 | -0.032 | -0.424*** |
| H3K27me3 | -0.080*** | 0.039 | 0.027 | -0.199*** |
| H3K27ac | -0.134*** | 0.10*** | 0.070** | -0.396*** |
| Transcription rate | -0.122*** | -0.033 | -0.043* | -0.214*** |
| H3K4me1PB | -0.119*** | 0.105*** | 0.080*** | -0.385*** |
| H3K9me3PB | 0.106*** | -0.169*** | -0.133*** | 0.420*** |
| Nucleosome occupancy | -0.070** | 0.224*** | 0.184*** | -0.357*** |
| DNAse hypersensitivity | -0.144*** | 0.086*** | 0.062** | -0.302*** |
| Replication time | -0.154*** | 0.045* | 0.019 | -0.474*** |
| GC content | -0.110*** | 0.167*** | 0.132*** | -0.324*** |

Many of the genomic variables are correlated to each other. If we use principle components to reduce the dimensionality, the first principle component (PC) explains 58% of the variation, the second 13%, the third and fourth 6.9 and 5.7% of the variation. We find that the density of DNMs is significantly correlated to the first PC in both datasets, but the correlation is negative for the Francioli data (r = -0.14, p<0.001) and positive for Wong (r = 0.14, p<0.001). Both are significantly positive correlated to the second PC (Francioli, r = 0.14, p<0.001; Wong, r = 0.27, p<0.001), uncorrelated to the third component and Wong is significantly correlated to the fourth component (r -0.059, p = 0.005).

It is possible that the differences between the Wong and Francioli datasets are due to biases in the ability to call DNMs. However, analysing the Wong data using the number of callable trios at each site does not qualitatively alter the pattern of correlation in the Wong dataset (Table 4) or the correlations to the principle components of the genomic features (PC1, r=0.11 p<0.001; PC2, r=0.25, p<0.001; PC3, r=-0.019, p=0.37; PC4, r=-0.048, p =0.019).

To investigate whether these patterns are consistent across mutational types, we calculated the correlation between the density of each mutational type (e.g. CpG C>T mutations at CpG sites) and the first two PCs of the genomic features. For the Francioli data the patterns are perfectly consistent; all mutational types, if they show a significant correlation, are significantly negatively correlated to the first PC, and significantly positively correlated to the second (Table S6). For the Wong data the patterns are more heterogeneous; all mutational types are positively correlated to the second PC, but some mutational types are significantly positively correlated to the first PC and others negatively correlated.

In order to try and disentangle which factors might be most important in determining the rate of mutation we used stepwise regression. We find, as expected, that the models selected for the Francioli and Wong datasets have some commonalities but some differences as well (Table 5); in both cases the density of DNMs is correlated to female recombination rate and negatively correlated to the density of H3K4me3. However, they are both correlated to DNAse hypersensitivity but in opposite directions and they have their own unique signatures; for example, in Francioli the density of DNMs is correlated to replication time whereas it is correlated to nucleosome occupancy in the Wong data. The differences are not due to variation in the ability to call DNMs in the Wong dataset since repeating the analyses using the sum of callable trios rather than sites, does not alter the patterns (Table 5). Intriguingly the regression models fitted to the Francioli and Wong data are quite similar at the 100KB scale (Table S7) despite the fact that the Francioli data is significantly negatively correlated to the first principle component, whereas Wong is significantly positively correlated.

**Table 5.** The standardised regression coefficients from a stepwise multiple regression with forward variable selection (parameter has to be significant at p<0.05 to be added to the model).

|  | Francioli | Wong | Wong<br>(callable) |
| --- | --- | --- | --- |
| Male recombination rate |  | 0.10*** | 0.10*** |
| Female recombination rate | 0.069** | 0.091** | 0.084** |
| H3K4me1 | 0.12** |  |  |
| H3K4me3 | -0.084* | -0.13*** | -0.13*** |
| H3K27me3 |  |  |  |
| H3K27ac |  |  |  |
| Transcription rate |  |  |  |
| H3K4me1PB |  |  |  |
| H3K9me3PB |  |  |  |
| Nucleosome occupancy |  | 0.49*** | 0.44*** |
| DNAse hypersensitivity | -0.12** | 0.14* | 0.15* |
| Replication time | -0.12** |  |  |
| GC content |  | -0.39** | -0.38** |
| $r^2$ | 0.044 | 0.10 | 0.084 |

The amount of variation explained by the multiple regression models is small – 0.044 and 0.10 for Francioli and Wong respectively - but this might be expected given the small number of DNMs per MB and hence the large sampling error. To investigate how much of the explainable variance the model explains we sampled rates from the gamma distribution fitted to the distribution of DNMs across the genome and generated DNMs using these rates and then correlated these simulated rates to the true rates (i.e. those sampled from the gamma distribution). The average coefficient of determination for the simulated data is 0.12 and 0.24 for the Francioli and Wong datasets respectively suggesting that the regression model explains ~37% and ~42% of the explainable variance for the two datasets. In both cases none of the simulated datasets have a coefficient of determination that is as low as the observed.

### Correlation with divergence

The rate of divergence between species is expected to depend, at least in part, on the rate of mutation. To investigate whether variation in the rate of substitution is correlated to variation in the rate of mutation we calculated the divergence between humans and chimpanzees, initially by simply counting the numbers of differences between the two species. There are at least three different sets of human-chimpanzee alignments: pairwise alignments between human and chimpanzee (PW)[41] found on the University of California Santa Cruz (UCSC) Genome Browser, the human-chimp alignment from the multiple alignment of 46 mammals (MZ)[42] from the same location, and the human-chimp alignment from the Ensembl Enredo, Pecan and Ortheus primate multiple alignment (EPO) [43]. We find that the correlation depends upon the human-chimpanzee alignments used and the amount of each window (either 1MB or 100KB) covered by aligned bases (Figure 5). The correlation is significantly negative if we include all windows for the UCSC PW and MZ alignments at the 1MB scale (similar results are obtained at 100KB), but becomes more positive as we restrict the analysis to windows with more aligned bases. In contrast, the correlations are always positive when using the EPO alignments, and the strength of this correlation does not change once we get above 200,000 aligned bases per 1MB. Further analysis suggests there are some problems with the PW and MZ alignments because divergence per MB window is inversely correlated to mean alignment length (r = -0.31, p < 0.0001) for the PW alignments and positively correlated (r = 0.57, p < 0.0001) for the MZ alignments (Figure S4). The EPO alignment method shows no such bias and we consider these alignments to be the best of those available. Therefore, we use the EPO alignments for the rest of this analysis.

**Figure 5.**
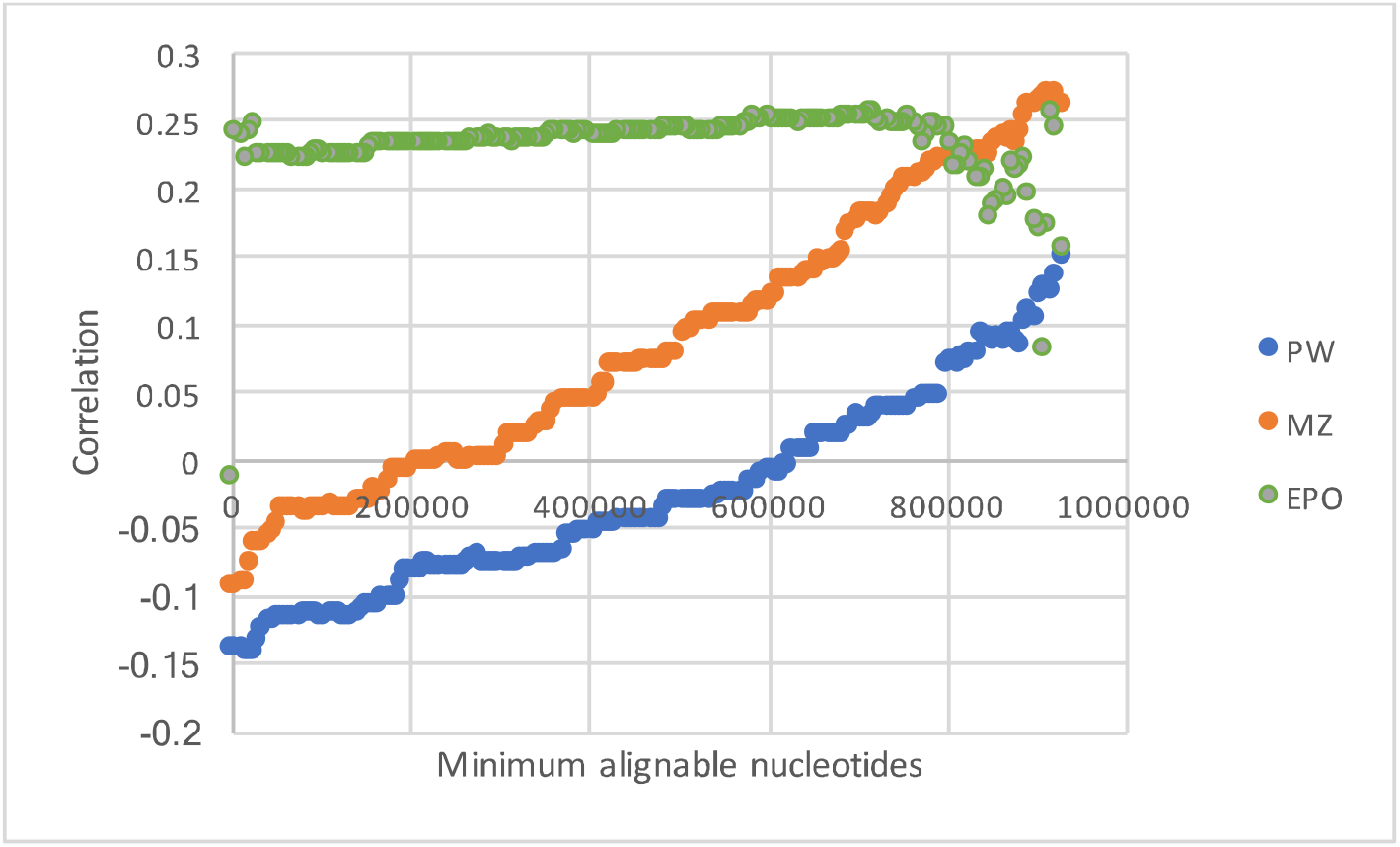
The correlation between the divergence from human to chimpanzee and the density of DNMs in humans as a function of the number of aligned sites per window for three sets of alignments: UCSC pairwise alignments (PW, blue), UCSC multi-way aligments (MZ, orange) and EPO multi-species alignments alignments (EPO, green).

To gain a more precise estimation of the number of substitutions we used the method of Duret and Arndt [21], which is a non-stationary model of nucleotide substitution that allows the rate of transition at CpG dinucleotides to differ to than that at other sites. As expected the divergence along the human lineage (since humans split from chimpanzees) is significantly correlated to the rate of DNMs (Francioli = 0.21 p<0.001; Wong = 0.11, p<0.001). However, the correlation between the rate of DNMs and divergence is not expected to be perfect even if variation in the mutation rate is the only factor affecting the rate of substitution between species; this is because we have relatively few DNMs and hence our estimate of the density of DNMs is subject to a large amount of sampling error. To investigate how strong the correlation could be, we follow the procedure suggested by Francioli et al. [3]; we assume that variation in the mutation rate is the only factor affecting the variation in the substitution rate across the genome between species and that we know the substitution rate without error (this is an approximation, but the sampling error associated with the substitution rate is small relative to the sampling error associated with DNM density because we have so many substitutions). We generated the observed number DNMs according to the rates of substitution, and then considered the correlation between these simulated DNM densities and the observed substitution rates. We repeated this procedure 1000 times to generate a distribution of expected correlations. Performing this simulation, we find that we would expect the correlation between divergence and DNM density to be 0.33 and 0.48 for the Francioli and Wong datasets respectively, considerably greater than the observed values of 0.21 and 0.11 respectively. In none of the simulations was the simulated correlation as low as the observed correlation.

There are several potential explanations for why the correlation is weaker than it could be; the pattern of mutation might have changed [37, 44–46], or there might be other factors that affect divergence. Francioli et al. [3] showed that including recombination in a regression model between divergence and DNM density significantly improved the coefficient of determination of the model; a result we confirm here; the coefficient of determination when recombination is included in a regression of divergence versus DNM density increases from 0.042 to 0.074 and from 0.013 to 0.041 for the Francioli and Wong datasets respectively.

As detailed in the introduction there are at least four explanations for why recombination might be correlated to the rate of divergence independent of its effect on the rate of DNM: (i) biased gene conversion, (ii) recombination affecting the efficiency of selection, (iii) recombination affecting the depth of the genealogy in the human-chimpanzee ancestor and (iv) problems with regressing against correlated variables that are subject to sampling error. We can potentially differentiate between these four explanations by comparing the slope of the regression between the rate of substitution and the recombination rate (RR), and the rate DNM and the RR. If recombination affects the substitution rate, independent of its effects on DNM mutations, because of GC-biased gene conversion (gBGC), then we expect the slope between divergence and the RR to be greater than the slope between DNM density and the RR for Weak>Strong (W>S), smaller for S>W, and unaffected for S<>S and W<>W changes. The reason is as follows; gBGC increases the probability that a W>S mutation will get fixed but decreases the probability that a S>W mutation will get fixed. This means that regions of the genome with high rates of recombination will tend to have higher substitution rates of W>S mutations than regions with low rates of recombination hence increasing the slope of the relationship between divergence and recombination rate. The opposite is true for S>W mutations, and S<>S and W<>W mutations should be unaffected by gBGC. If selection is the reason that divergence is correlated to recombination independently of its effects on the mutation rate, then we expect all the slopes associated with substitutions to be less than those associated with DNMs. The reason is as follows; if a proportion of mutations are slightly deleterious then those will have a greater chance of being fixed in regions of low recombination than high recombination. If the effect of recombination on the substitution rate is due to variation in the coalescence time in the human-chimp ancestor, then we expect all the slopes associated with substitution to be greater than those associated with DNMs; this is because the average time to coalescence is expected to be shorter in regions of low recombination than in regions of high recombination. Finally, if the effect is due to problems with multiple regression then we might expect all the slopes to become shallower. Since the DNM density and divergences are on different scales we divided each by their mean to normalise them and hence make the slopes comparable.

The results of our test are consistent with the gBGC hypothesis; the slope of divergence versus RR is greater than the slope for DNM density versus RR for W>S mutations and less for S>W mutations (Figure 6); we present the analyses using sex-averaged RR, but the results are similar for either male or female recombination rates, and for 100KB windows (Figures S5 and S6 and Tables S8 and S9). These differences are significant in the expected direction for all comparisons except W>S from the Wong data (Table 6)(significance was assessed by bootstrapping the data by MB regions 100 times and then recalculating the slopes). There are no significant differences between the slope for W<>W and S<>S mutations and the slope for substitutions, again a result consistent with gBGC.

**Figure 6.**
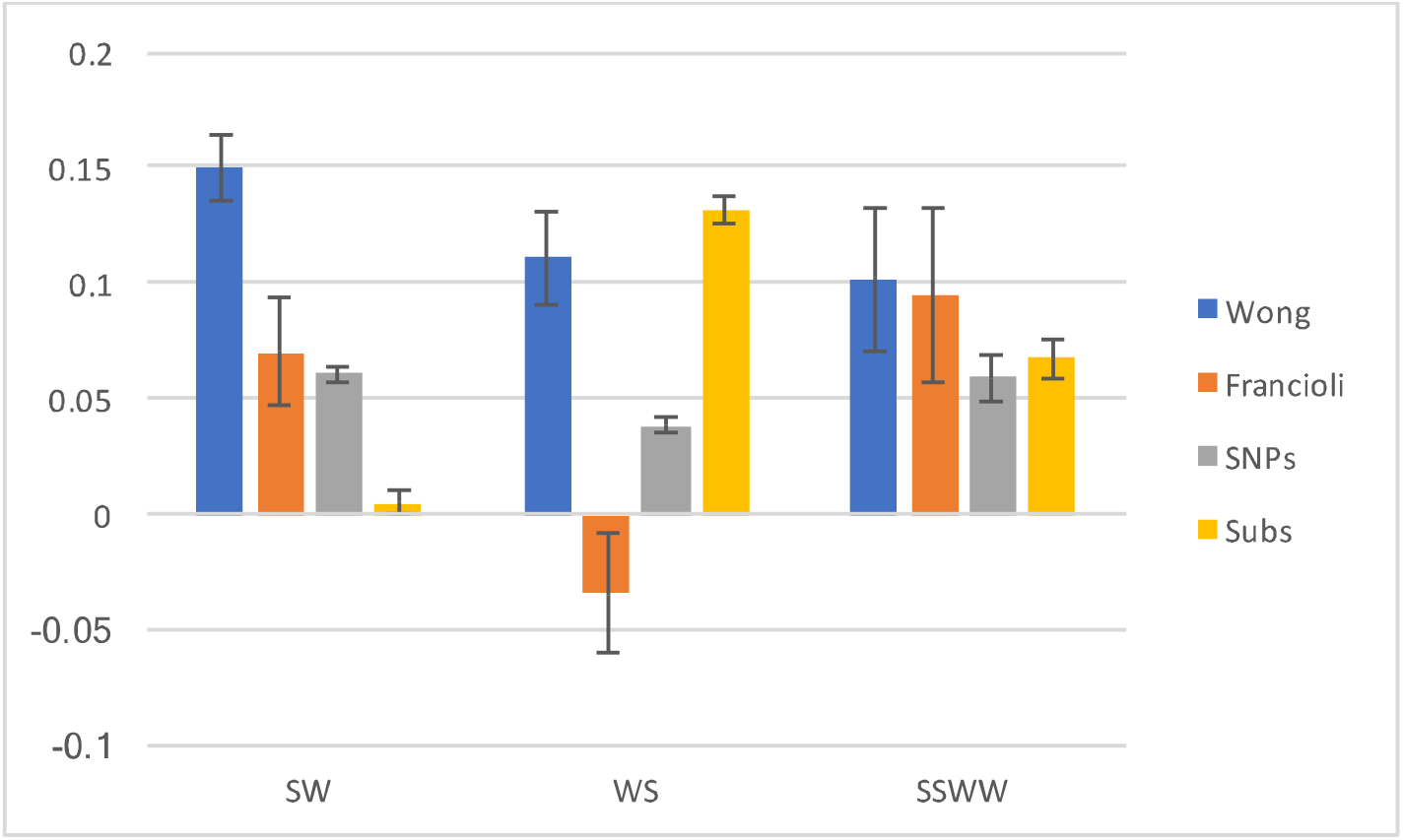
The slope (and SE) between normalised DNM density and normalised sex-averaged recombination rate (RR) (Wong - blue, Francioli - orange), normalised SNP density and RR (grey), and normalised substitution density and RR (yellow). In each case the values were normalised by dividing by the mean.

**Table 6.** Proportion of bootstrap replicates in which the slope of the normalised DNM density versus sex-averaged recombination rate, is greater than the slope of the normalised number of substitutions (or SNPs) versus recombination rate. 100 bootstrap replicates were performed in each case.

|  | SW | WS | SSWW |
| --- | --- | --- | --- |
| Wong v Subs | 1 | 0.17 | 0.89 |
| Francioli v Subs | 1 | 0 | 0.81 |
| Wong v SNPs | 1 | 1 | 0.95 |
| Francioli v SNPs | 0.66 | 0 | 0.76 |

### Correlation with diversity

Just as we expect there to be correlation between divergence and DNM rate, so we might expect there to be correlation between DNA sequence diversity within the human species and the rate of DNM. To investigate this, we compiled the number of SNPs in 1MB and 100KB windows from the 1000 genome project [47, 48]. There is a positive correlation between SNP density and DNM rate in both datasets (Francioli r = 0.19 p<0.001; Wong r = 0.30, p<0.001).

Using a similar strategy to that used in the analysis of divergence we calculated the correlation we would expect if all the variation in diversity was due to variation in the mutation rate by assuming that the level of diversity was known without error, and hence was a perfect measure of the mutation rate (we have on average 31,000 SNPs per MB, so there is little sampling error associated with the SNPs). We then simulated the observed number of DNMs according to these inferred mutation rates. The expected correlations are 0.22 and 0.32 in the Francioli and Wong datasets; these are not significantly greater than the observed correlation (p=0.1 in both cases). A similar pattern is observed for individual mutational types at both the 1MB and 100KB scale, with some being greater and others smaller than expected (Table S10). These results suggest that the majority of the variation in diversity at the 1MB scale is due to variation in the mutation rate.

Although much of the variation in diversity appears to be due to variation in the mutation rate we tested for the effect of gBGC. We find that the slope of SNP density versus RR is lower than the slope between DNM density and RR for S>W mutations, greater for W>S mutations and not significantly different for S<>S/W<>W mutations for the Francioli dataset. Furthermore, the slope between SNP density and RR is between the slopes for DNM density versus RR and substitutions and RR. This is consistent with gBGC, since gBGC is expected to have smaller effects on diversity than divergence. However, in the Wong dataset the DNM slope is significantly greater than the diversity slope for all mutational types consistent with a direct effect of selection on diversity.

### Divergence to other species

The divergence between species, usually humans and macaques, is often used to control for mutation rate variation in various analyses. But how does the correlation between divergence and the DNM rate in humans change as the species being compared get further apart? Terekhanova et al. [46] showed that the rate of S<>S and W<>W substitutions (chosen to eliminate the influence of gBGC) along the human lineage at the 1MB scale is correlated to that along other primate lineages, but that the correlation declines as the evolutionary distance increases. This suggests that the mutation rate evolves at the 1MB relatively rapidly. However, they did not consider DNMs in detail. To investigate further, we compiled data from a variety of primate species – human/chimpanzee/orang-utan (HCO) considering the divergence along the human and chimp lineages, human/orangutan/macaque (HOM) considering the divergence along the human and orangutan lineages, and human/macaque/marmoset (HMM) considering the divergence along the human and macaque lineages. This yields two series of divergences of increasing evolutionary divergence: the human lineage from HCO, HOM and HMM, and chimp from HCO, orangutan from HOM and macaque from HMM. We estimated the divergence using the non-stationary method of Duret and Arndt [21] that treats CpG sites separately. We do not restrict ourselves only to DNMs in the aligned regions but used all DNMs in each window. In this way, the average number of DNMs per window is independent of the evolutionary divergence. As expected, we find that the correlation between the density of DNM and the rate of substitution declines as the evolutionary divergence increases (Figure 7).

**Figure 7.**
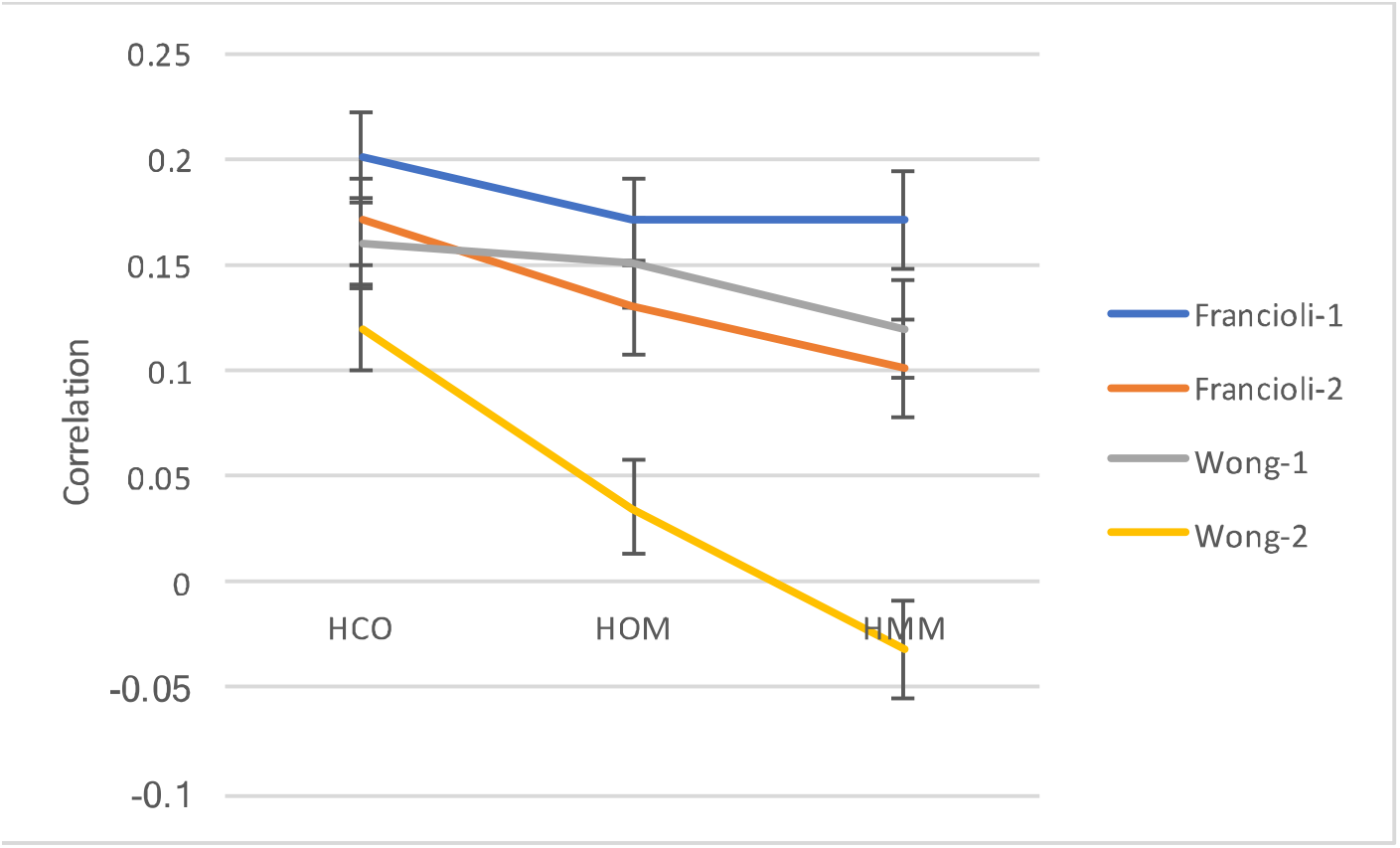
The correlation between DNM density and the substitution rate for different branches. Francioli-1 and Wong-1 are the correlations involving the divergence along the human lineages from the comparison of human-chimp-orangutan (HCO), human-orangutan-macaque (HOM) and human-macaque-marmoset (HMM). Francioli-2 and Wong-2 involve the divergences along the chimpanzee, orangutan and macaque lineages

## Discussion

We have considered the large-scale distribution of DNMs along the human genome using a meta-analysis of 4 datasets of DNMs obtained by the sequencing of trios (an individual and their parents). Unfortunately, there are significant differences between these datasets: they differ in the amount of variation they show and the two largest datasets, which we have studied in detail, show some conspicuous differences; in particular the DNMs from the studies of Francioli et al. [3] and Wong et al. [6] correlate differently to genomic variables such as replication time. Patterns that exist in both datasets likely reflect true patterns associated with DNMs. Patterns that differ between datasets may be real and reflect differences in age, environment and ethnicity of the individuals sequenced in each study, or they may be due to systematic biases in the discovery of DNMs.

The patterns and results that are common to both datasets are as follows. There appears to be rather little variation in the mutation rate at a large scale, however, there is variation at a large scale that cannot be explained by variation at smaller scales, and large-scale variation forms part of a continuum. Furthermore, we can conclude that the level of variation for different mutational types is similar and different mutational types covary together. We find no evidence that there is variation in the pattern of mutation that generates variation in GC content. We confirm the correlation between the mutation rate, as measured by DNM density, is not as strong as it could be, and demonstrate that this is in part due to BGC. In contrast, we find that variation in diversity at large scales is largely a consequence of variation in the mutation rate. Finally, we demonstrate that the correlation between the rate of DNM and the rate of substitution, declines as increasingly divergent species are considered.

Whether the differences between the datasets are due to error or differences between the sampled individuals is not clear. it is possible that some of the differences between the Francioli and Wong datasets are a product of the sequencing technology and processing used to call DNMs. It has been noted before that the pattern of mutation at the single nucleotide level varies [35, 36], and the rate at which the mutation rate increases with paternal age [36] differs between studies; observations which have been interpreted as being artefcats of sequencing or processing [35, 36]. However, these differences and those reported here between the Francioli and Wong datasets could be due to differences in age, environment or ethnicity. it has been shown that mutations born to younger fathers are enriched in late replicating DNA, whereas this pattern is not observed in older fathers [3]. Thus at least part of the difference between the Francioli and Wong datasets could be due to the paternal age. The average age of fathers in the Wong dataset was 33.4 years [6], in the Francioli dataset it was 27.7 years (Francioli pers comm). We might therefore not expect to see such a strong replication time effect in the Wong data. Since replication time is strongly correlated to many of the other genomic variables - it is highly correlated to the first PC of the genomic variables r = 0.80, p<0.001 - this one effect might explain why the patterns of correlation are different in the two datasets.

If paternal age is the reason for the difference between the Wong and Francioli datasets then this implies that the majority of mutations over human evolution have come from relatively young fathers since the rate of S<>S and W<>W substitutions (chosen to remove the effects of gBGC) show similar correlations to genomic variables as the Francioli DNM data (Table 4 and Table S7 for 100KB).

It is also possible that the differences between the Francioli and Wong datasets are due to ethnicity. It has recently been shown that the rate and pattern of mutation varies over very short timescales such that they can vary between human populations [37, 44, 45]; for example the rate of TCC to TTC is elevated in Europeans [37, 44]. At larger scales Terekhanova et al. [46] considered the correlation between the S<>S and W<>W substitution rate along the human lineage and other primate lineages at the 1MB scale. They showed that the strength of the correlation declined as more distant species were considered suggesting that the mutation rate evolves at this scale as well as at finer scales. However, the rate of decline was fairly slow, and human populations would not be predicted to show very different patterns from this analysis.

The evolution of the large-scale variation in GC-content across the human genome has been the subject of much debate [25]. Mutation bias [14–18], selection [13, 19, 20] and biased gene conversion [21–24] have all been proposed as explanations. There is good evidence that biased gene conversion has some effect on the base composition of the human genome [24, 26–28]. However, this does not preclude a role for mutation bias. We have tested the mutation bias hypothesis using the DNM data and two different tests. We find no evidence that the pattern of mutation varies across the genome in a way that would generate variation in GC-content. Instead we provide additional evidence that biased gene conversion influences the chance that mutations become fixed in the genome.

As expected the rate of divergence between species is correlated to the rate of DNM, however, the strength and even the sign of the correlation depends on the alignments being used. The correlations between divergence and DNM density are actually negative if no filtering is applied to the UCSC alignments, and there is a negative correlation between divergence and alignment length for the pairwise alignments from the UCSC genome browser, and a positive correlation for the multi-species alignment. It is clear that there are problems with these alignments and that they should be used with caution.

As Francioli et al. [3] showed, the correlation between divergence and DNM density is worse than it would be if variation in the mutation rate was the only factor affecting divergence. This is perhaps not surprising because the substitution rate depends both on the rate of mutation and the probability of fixation, both of which may vary across the genome. Francioli et al. [3] further demonstrated that although the rate of DNM is correlated to the rate of recombination, divergence is correlated to the rate of recombination independently of this effect. There are at least four explanations for the effect of recombination on divergence: (i) biased gene conversion, (ii) direct selection, (iii) linked selection and (iv) problems with multiple regression. We have provided evidence for an effect of biased gene conversion, but no obvious influence of three other factors – i.e. the slope of the regression between DNM density and RR is not significantly different to the slope of the regression between divergence and RR for S<>S and W<>W mutations. The fact that there is no obvious effect of indirect selection is surprising given the results of McVicker et al. [29]. They showed that the divergence between humans and chimpanzees was significantly lower near exons and other regions of the genome subject to evolutionary constraint. A similar reduction was not observed in the divergence of human and macaque and human and dog, suggesting that the pattern was not due to selection outside exons, or regions identified as being subject to selection. They therefore inferred that the reduction was due to the effect of linked selection reducing diversity in the human-chimpanzee ancestor. There are several possible reasons why see no evidence of this effect in our analysis. First, our test may not be powerful enough. Second, the effects may be counteracted by direct selection which is expected to affect the slope of the regression between divergence and RR in the opposite direction to indirect selection. Third, the scale, magnitude and variation in the effects of indirect selection may be not large enough to affect the relationship between divergence and the rate of mutation; if there is little variation in the magnitude of the indirect effects of selection across the genome at the 1MB (or 100KB) level then indirect selection will have no effect on the correlation between the rate of mutation and divergence.

In contrast to the pattern with divergence, we find that most of the variation in diversity, at least at the 1MB and 100KB scales, can be explained by variation in the mutation rate. Considering this, it is perhaps not surprising that the analysis of DNM density versus RR and SNP density versus RR slopes are inconclusive. The results from Francioli are consistent with BGC affecting he relationship between SNP and DNM density, but the data from Wong are consistent with a direct effect of selection on diversity. Again we find no evidence of an effect of indirect selection contrary to previous analyses [29, 49]. This may be because our analysis lacks power, or that the indirect effects of selection do not vary sufficiently at the scale we have analysed.

Divergence between species has often been used to control for mutation rate variation in humans (for example [29, 49’51]). This is clearly not satisfactory given that the correlation between divergence and rate of DNM is only about half as strong as it could and this correlation gets worse as more divergent species are considered. Unfortunately, correcting for mutation rate variation is likely to be difficult because attempting to predict mutation rates from genomic features is unreliable, given the failure of the regression analyses to explain more than half the explainable variation. Furthermore, the largest amounts of variation are at the smallest scales (Figure 3B) where we have the lowest density of DNMs.

It has been known for sometime, that diversity across the human genome is correlated to the rate of recombination[30-32] and there has been much debate about whether this is due to mutagenic effects of recombination or the effect of recombination on processes such as genetic hitch-hiking and background selection. Divergence between humans and other primates is correlated to the rate of recombination, which was initially interpreted as being due to a mutagenic effect of recombination [30, 32] but subsequently it has been interpreted as evidence of gBGC [21]. Both of these hypotheses appear to be correct – the rate of DNM is correlated to the rate of recombination ([3]; results above), but recombination also affects which mutations become fixed through gBGC.

We find, as others have before [3, 5], that the rate of germ-line DNM is correlated to a number of genomic features. However, we find that these features explain less than 50% of the explainable variance leaving the majority of the variance unexplained. Our inability to predict the mutation rate might be because the genomic features have not been assayed in the relevant tissue, the germ-line, or that there are important features that have yet to be assayed. Interestingly, Terekhanova et al. [46] showed that this unexplained component of the substitution rate evolves more rapidly than the explained component. They demonstrated that the substitution rates at the 1MB level in a range of primate species was almost as well correlated to genomic features in humans, as the substitution rate along the human lineage. This implies that the variance in the substitution rate not explained by genomic features, evolves rapidly, given that the correlation between the substitution rate in humans and other lineages declines as they get more distant, There is clearly much we do not currently understand about the why there is large scale variation in the mutation rate and how it evolves through time.

## Materials and methods

### DNM data

Details of DNM mutations were downloaded from the supplementary tables of the respective papers: 26,939 mutations from Wong et al. [6], 11016 mutations from Francioli et al. [3], 4931 mutations from Kong et al. [4] and 547 mutations from Michaelson et al. [5]. These were all mapped to hg19/GRCh37. Only autosomal DNMs were used.

### Alignments

Three sets of alignments were used in this analysis, all based on human genome build hg19/GRCh37: (i) the University of California Santa Cruz (UCSC) pairwise (PW) alignments [41] for human-chimpanzee (hg19-panTro4 downloaded from http://hgdownload.cse.ucsc.edu/goldenpath/hg19/vsPanTro4/) (ii) the UCSC MultiZ (MZ) 46-way alignments [42] downloaded from http://hgdownload.cse.ucsc.edu/goldenpath/hg19/multiz46way/ and (iii) Ensembl Enredo, Pecan, Ortheus (EPO) 6 primate multiple alignment, release 74, [43] downloaded from ftp://ftp.ensembl.org/pub/release-74/emf/ensembl-compara/epo_6_primate/. We found that the EPO alignments were the most reliable – see main text – and they were used for the majority of the analyses.

### Selection and filtering of SNPs

All SNPs from the 1000 genomes project phase 3 [48] were downloaded from hgdownload.cse.ucsc.edu/gbdb/hg19/1000Genomes/phase3/. After removing all multi-allelic SNPs and, structural variants and indels we were left with 77,818,368 autosomal SNPs. After filtering out windows which had less than 50% of nucleotides aligning between human-chimpanzee-orangutan and no recombination rate scores we were left with 71,917,321 SNPs.

### Mutational models

We considered how well the variation at the 100KB and 1MB scale was predicted by two models of mutation rates: the rates estimated by Aggarwala et al. [38] based on the 7-mer context surrounding a site, and the rates estimated for each site by Michaelson et al. based on a variety of genomic features. The rates for Aggarwala et al. [38] were taken from their supplementary table 7, and the context of each site was used to predict the average mutation rate for each 100KB or 1MB window using their model. The mutability indices from the Michaelson et al. study [5] were provided by the authors. The analysis of the model of Michaelson et al. [5] is more complex since they give the probability of detecting a DNM in their data at each site in the genome, referred to as the mutability index (MI), but these do not translate directly into mutation rates. Using their DNM data we tabulated the number of sites in the genome with a given MI along with the number of DNMs from their study that had been observed at those sites. Because DNMs are not observed at some MIs we grouped MIs into groups of ten starting from the first MI with at least one DNM. We then regressed the log of the number of DNMs over the number of sites against the mean MI (see Figure S7). The regression line was estimated to be log(mutation rate) = -6.73 + 0.0103 x MI. Using this equation, we predicted the mutation rate at each site in the genome. Michaelson et al. [5] give MIs mapped to hg18; we lifted these over the hg19 using the liftover tool.

### Genomic features

Male, female and sex-averaged standardised recombination rate data [52] were downloaded from http://www.decode.com/additional/male.rmap, which provides recombination rates in 10KB steps. For each 100KB and 1MB windows the recombination rate was calculated as the mean of these scores with a score assigned to the window in which the position of its first base resided. GC content was calculated directly from the human genome (hg19/GCRh37) for 100kb and 1Mb windows. All other feature data was taken from the ENCODE project [53] and downloaded from the UCSC genome browser. Where possible we used data from the embryonic stem cell line H1- hesc. The mean value was taken for each genome feature across the window. For replication time data, we downloaded the ENCODE Repli-seq wavelet smoothed signal data [54, 55], provided in 1KB steps, for the GM12878, HeLa, HUVEC, K562, MCF-7 and HepG2 cell lines. Replication times were assigned to windows based upon their start coordinates. We computed the mean replication time for all autosomes for 100KB and 1Mb windows across all 6 cell lines. We measured transcription rate using RNA-seq data. Nucleosome occupancy was taken from the GM12878 cell line, histone modifications and RNA-seq data from the stem cell line H1-hesc. We only included windows in our analysis in which >50% of the window had data from all features.

### Statistical analysis

SPSS version 22 and Mathematica version 10 were used for all statistical analyses.

To estimate the mutation rate distribution we use the method of [8]. In brief we assume that the mutation rate in each window is 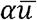 where 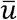 is the average mutation rate per site and α is the rate above or below this mean. α is assumed to be gamma distributed. The number of mutations per window is assumed to be Poisson distributed with a mean 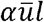 where *l* is the length of the window. This means that the number of mutations per window is a negative binomial. In considering a particular category of mutations, such as CpG transitions, we considered the number of CpG transition DNMs at CpG sites. We fit the distribution using maximum likelihood using the *NMaximize* function in *Mathematica*. Initial analyses suggested that the maximum likelihood value of the mutation rate parameter was very close to the mean estimate of the mutation rate; as a consequence to speed up the maximization we fixed the mutation rate to its estimated mean and found the ML estimate of the shape parameter of the gamma distribution.

We investigated the correlation between different types of mutation across windows by fitting a single distribution to both types of mutation, estimating the shape parameter of the shared distribution as the mean of the CV of the ML estimates of distributions fitted to the two categories independently. We then used this distribution to simulate data; we drew a random variate for each window from the distribution assigning this as the rate for that window. We then generated two Poisson variates with the appropriate means such that the total number of DNMs for each type of mutation was expected to be equal the total number of DNMs of those types.

To test whether the mutation pattern varied across the genome in a manner that would generate variation in the mutation rate we fit the following model. Let us assume that the mutation rate from strong (S) to weak (W) base pairs, where strong are G:C and weak are A:T, be *μ(1–f_e_)*, where μ is the mutation rate and *f_e_* is the equilibrium GC-content to which the sequence would evolve if there was no selection or biased gene conversion. Let the mutation rate in the opposite direction be *μf_e_* and the current GC-content be *f*. Then we expect the proportion of mutations that are S->W to be

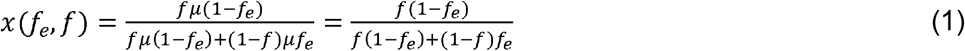

Let us assume that *f_e_* is normally distributed. Then the likelihood of observing *i* S>W mutations out of a total of *n* S>W and w>S mutations is

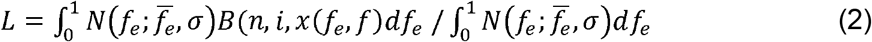

The total loglikelihood is therefore the sum of the log of equation 2 for each MB or 100KB window across all the windows in the genome. The maximum likelihood values were obtained by using the *NMaximize* routine in *Mathematica*.

### Simulations

In a number of analyses, we simulate DNMs under assumed model; for example, using the 7-mer model of Aggarwala et al. [38]. In these simulations, we calculate the expected number of DNMs given the window’s mutation rate, the number of relevant sites and the total number of DNMs, and then generated a random Poisson variate from this expectation. In each simulation, we generated 1000 simulated datasets.

## Data availability

The DNM data is available from the supplementary information of the original papers or from the corresponding authors: (i) Michaelson et al. [5] Supplementary table 1 at http://www.cell.com/cell/fulltext/S0092-8674(12)01404-3; (ii) Kong et al. [4] Supplementary data http://www.nature.com/nature/journal/v488/n7412/full/nature11396.html#supplementary-information; (iii) Francioli et al. [3] from Shamil Sunyaev; (iv) Wong et al. [6] from supplementary table 1 https://www.nature.com/articles/ncomms10486. The number of callable trios at each site was provided by Wendy Wong. The human genome assembly hg19 and all genomic features relating to the assembly were downloaded from the UCSC genome browser www.genome.ucsc.edu. The 1000 genome data were also downloaded there. The mutatibility indices from Michaelson et al. [5] were provided by Jake Michaelson. The rates for Aggarwala et al. [38] were taken from their supplementary table 7 http://www.nature.com/ng/journal/v48/n4/full/ng.3511.html.

## Acknowledgements

We thank Wendy Wong for providing data, in particular information on the number of trios that were callable at each site, to Shamil Sunyaev for helpful discussion about their results and analysis, and to anonymous referees for helpful suggestions.

## Supplementary tables

**Table S1.** The CV of the gamma distribution fitted to the Wong and Francioli datasets separately at the 100KB scale.

|  | Francioli 100KB | Wong 100KB |
| --- | --- | --- |
| Overall | 0.33 (0.26, 0.40) | 0.34 (0.32, 0.38) |
| CpG | 0.54 (0.14, 0.79) | 0.43 (0.27, 0.57) |
| nonCpG | 0.35 (0.26, 0.43) | 0.33 (0.30, 0.37) |
| CpG ts | 0.53 (0, 0.79) | 0.45 (0.27, 0.59) |
| CpG tv | 0.055 (0.05, 0.061) | 0.036 (0, 0.88) |
| nonCpG ts | 0.33 (0.15, 0.45) | 0.29 (0.23, 0.35) |
| nonCpG tv | 0.55 (0.38, 0.69) | 0.40 (0.32, 0.48) |
| S>W | 0.38 (0.23, 0.49) | 0.38 (0.32, 0.43) |
| W>S | 0.077 (0, 0.087) | 0.29 (0.18, 0.37) |

**Table S2.** The p-value from a likelihood ratio test comparing a model in which the two categories of mutation have separate distributions to where they share one.

| Test | Francioli<br>100KB | Wong<br>100KB | Francioli<br>1MB | Wong<br>1MB |
| --- | --- | --- | --- | --- |
| CpG v nonCpG | ns | ns | <0.001 | ns |
| CpG ts v tv | ns | ns | ns | ns |
| nonCpG ts v tv | 0.031 | 0.025 | ns | ns |
| S>W v W>S | ns | ns | 0.019 | ns |

**Table S3.**
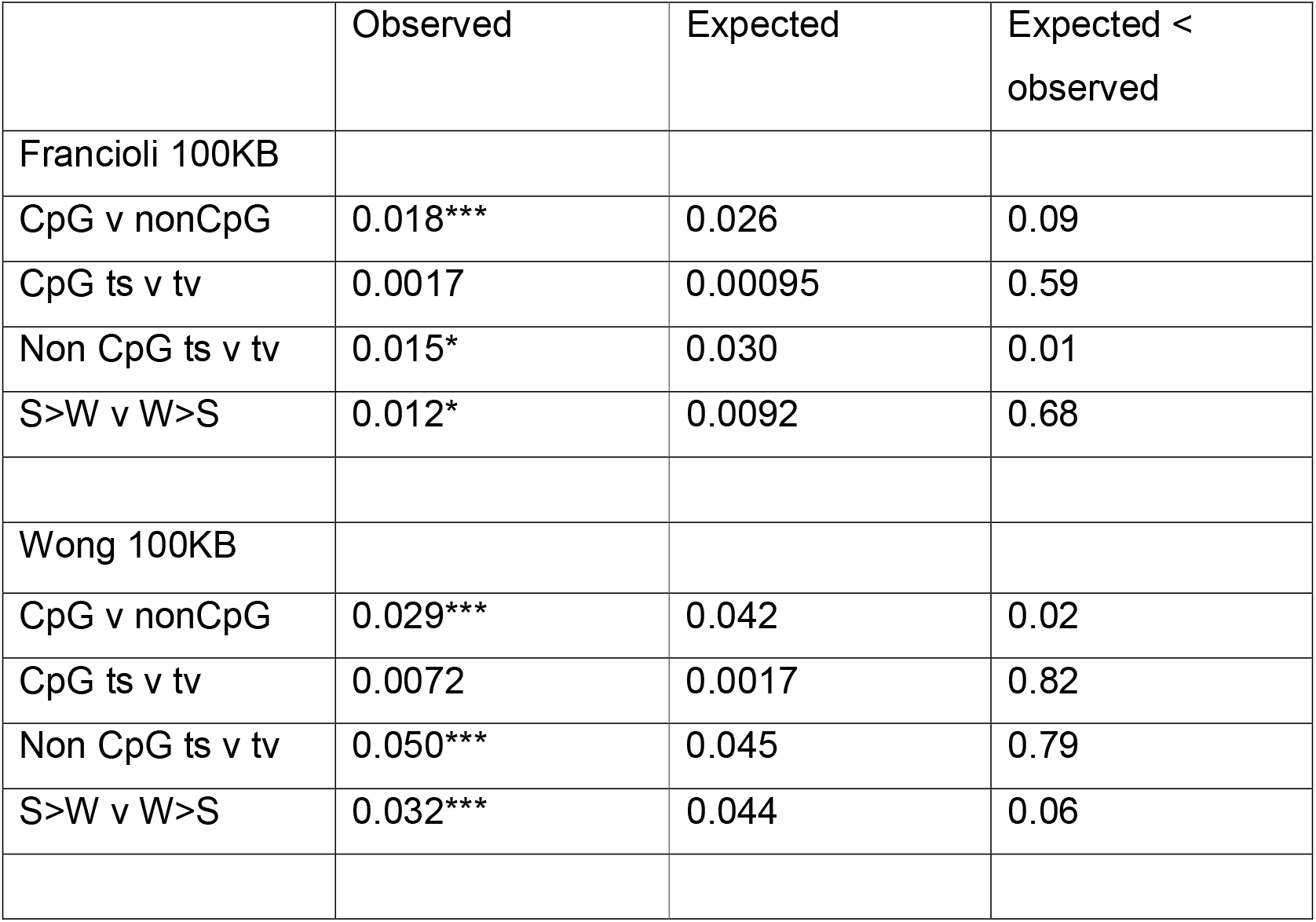
The correlation between different mutational types for the Francioli and Wong datasets separately at the 100KB scale.

**Table S4.** Correlation between the density of DNMs and the mutation rate estimates of Aggarwala et al. and Michaelson et al. at the 100KB scale. *p < 0.05 **p < 0.01 *** p < 0.001

|  | Aggarwala |  | Michaelson |  |
| --- | --- | --- | --- | --- |
|  | Observed | Simulated | Observed | Simulated |
| Francioli | -0.057*** | 0.060*** | 0.084*** | 0.41*** |
| Wong | 0.090*** | 0.094 | 0.018** | 0.57*** |

**Table S5.**
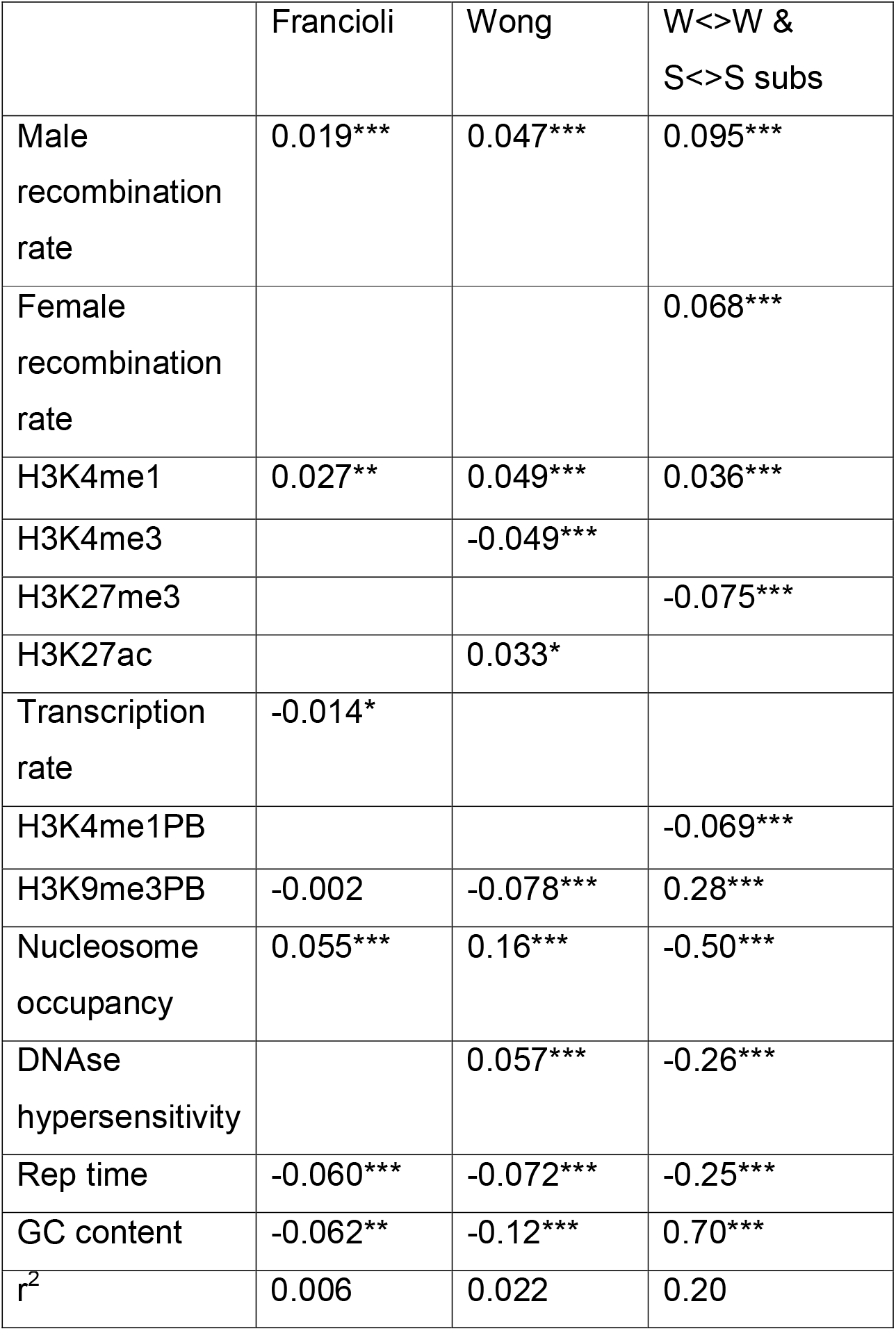
Correlations between the density of DNMs and various genomic features at the 100KB scale. * p<0.05, ** p<0.01, ***p<0.001

**Table S6.**
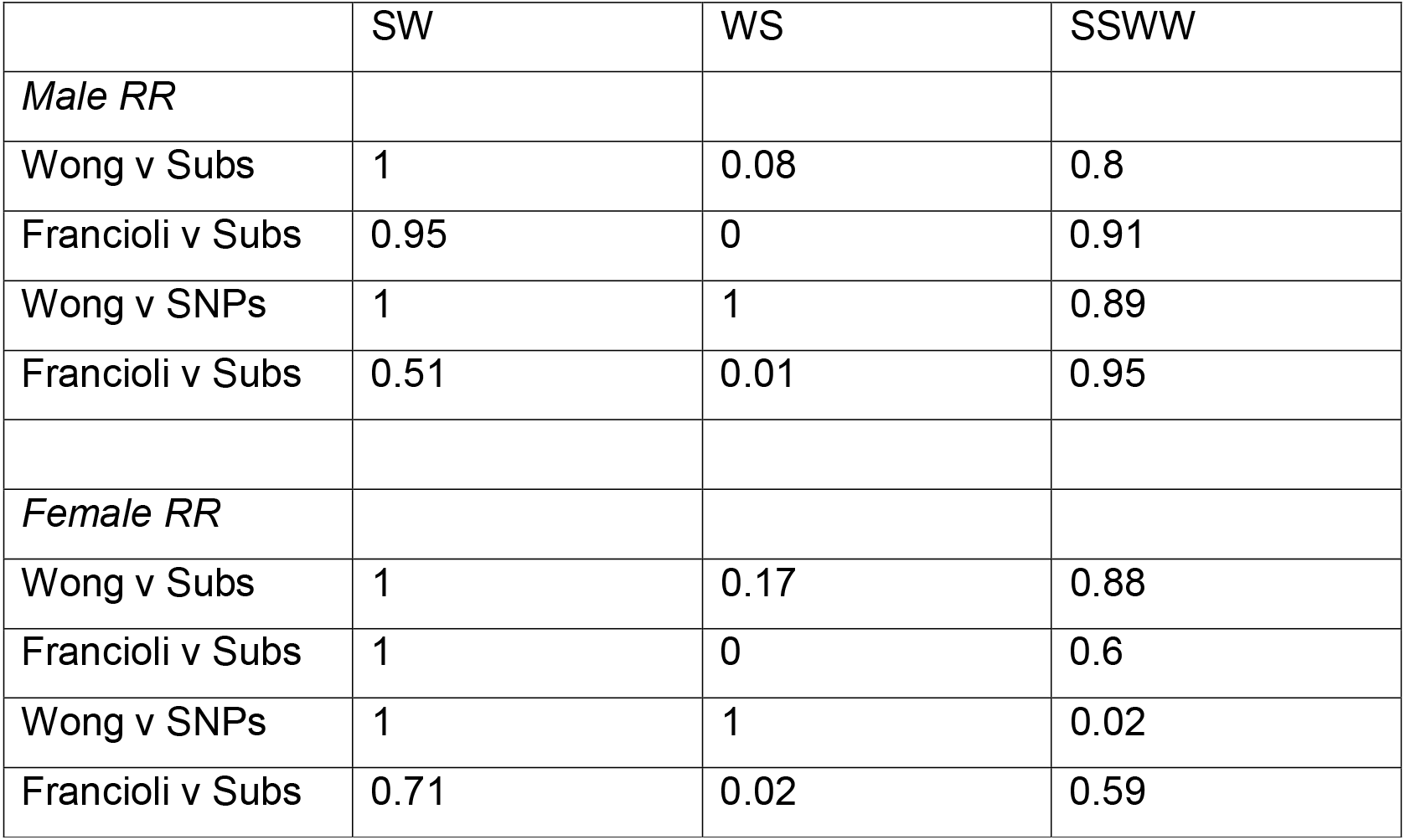
Correlations between the density of DNMs and the first two principle components of the genomic features. Note that DNM density is considered across the appropriate type of site (e.g. CpG C>T mutations at CpG sites). *p<0.05, **p<0.01, ***p<0.001.

**Table S7.**
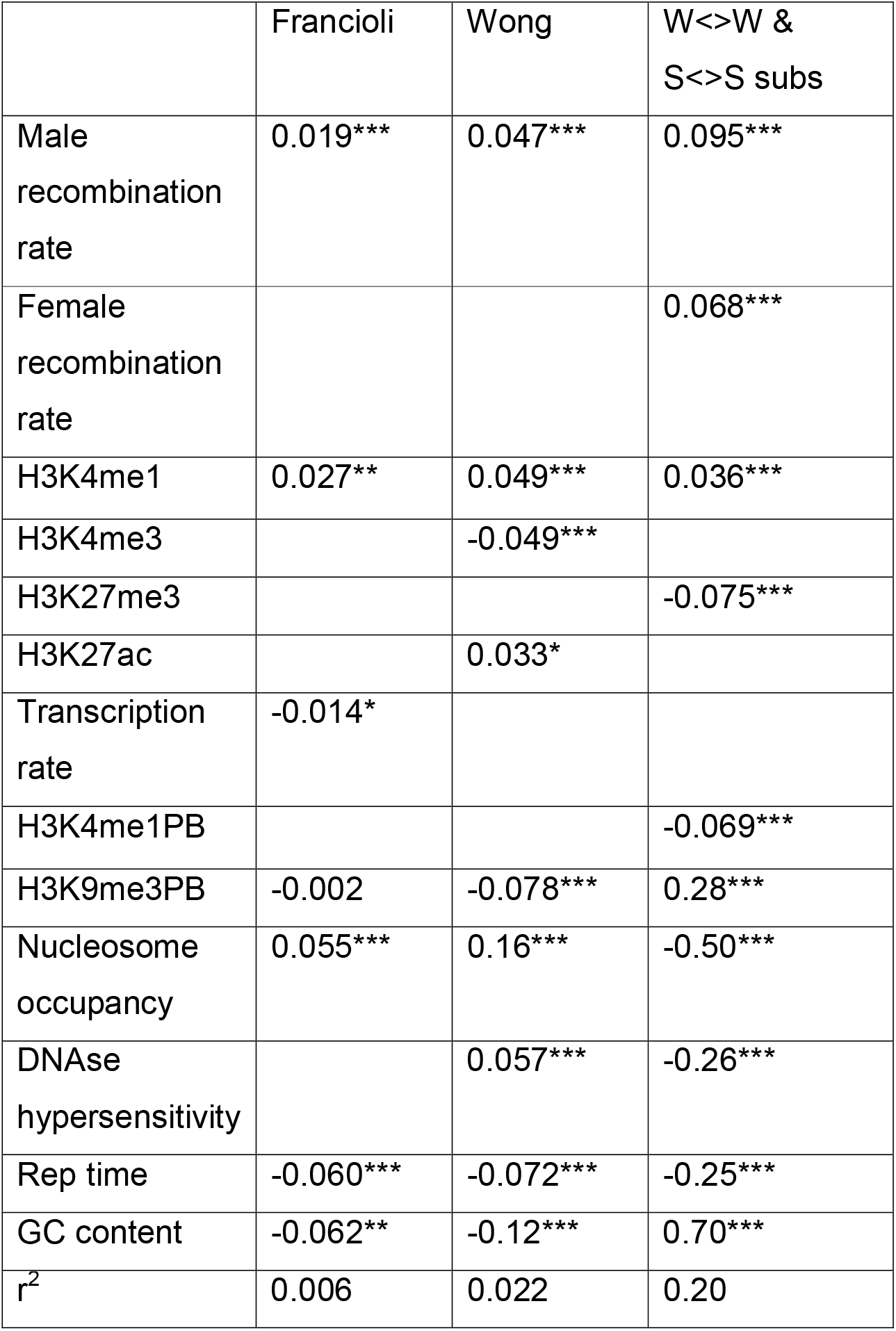
The standardised regression coefficients from a stepwise multiple regression with forward variable selection (parameter has to be significant at p<0.05 to be added to the model) at the 100KB scale.

**Table S8.**
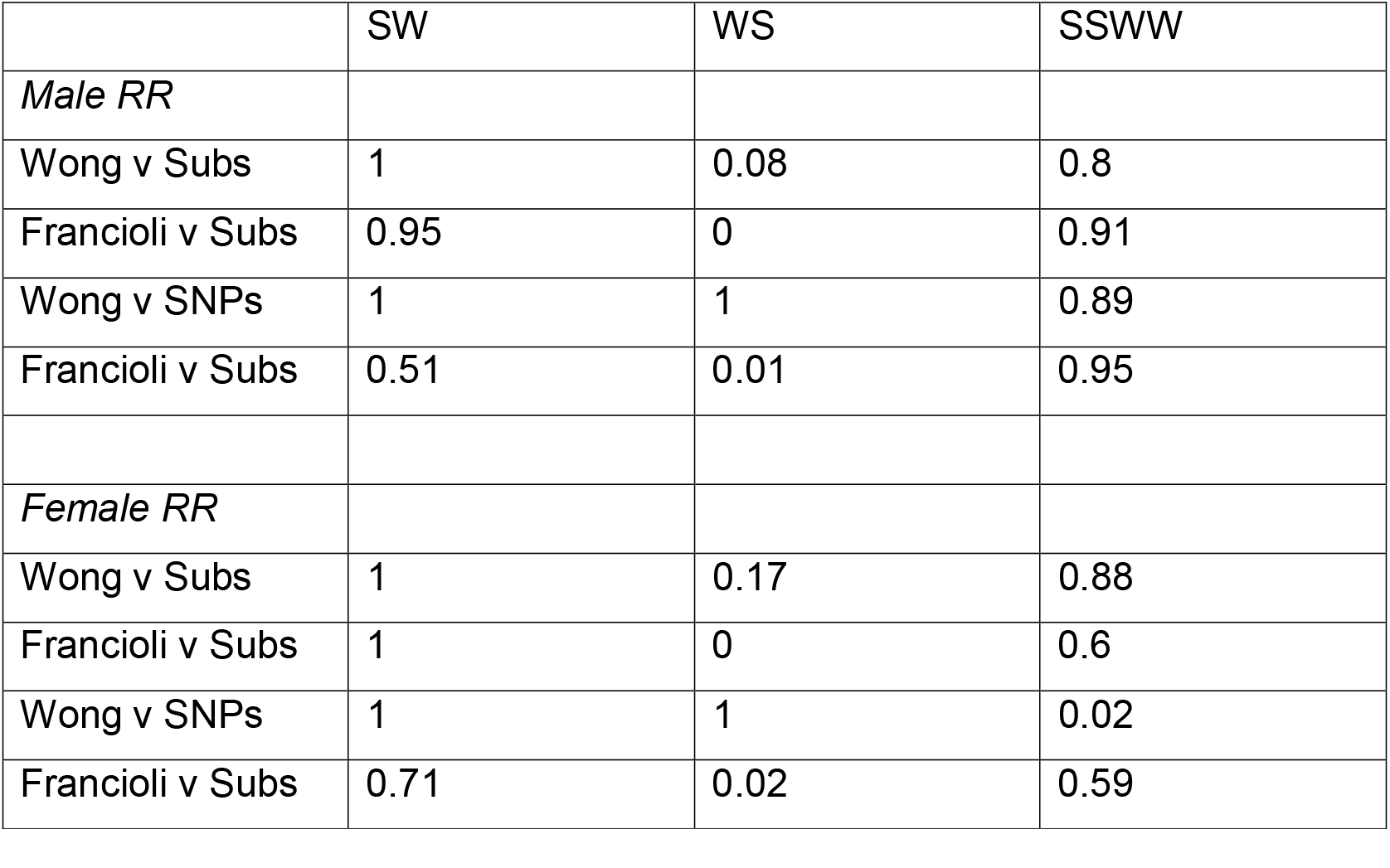
Proportion of bootstrap replicates in which the slope of the normalised DNM density at 1MB versus recombination rate, is greater than the slope of the normalised number of substitutions (or SNPs) versus recombination rate. 100 bootstrap replicates were performed in each case. Results are shown for male and female specific recombination rates.

**Table S9.** Proportion of bootstrap replicates in which the slope of the normalised DNM density at 100KB versus sex-averaged recombination rate, is greater than the slope of the normalised number of substitutions (or SNPs) versus recombination rate. 100 bootstrap replicates were performed in each case.

|  | SW | WS | SSWW |
| --- | --- | --- | --- |
| *Sex-averaged RR* |  |  |  |
| Wong v Subs | 1 | 0.11 | 1 |
| Francioli v Subs | 1 | 0 | 1 |
| Wong v SNPs | 1 | 1 | 1 |
| Francioli v SNPs | 0.84 | 0.02 | 0.61 |

**Table S10.** The observed and expected correlations between DNM and SNP density at the 1MB categories of mutations. The expected correlation is that expected if all the variation in SNP density is due to variation in the mutation rate; this was estimated by simulating data. *p<0.05, **p<0.01

|  | Francioli |  | Wong |  |
| --- | --- | --- | --- | --- |
|  | Observed | Expected | Observed | Expected |
| 1MB |  |  |  |  |
| All | 0.19** | 0.22 | 0.30** | 0.32 |
| CpG C>T | 0.20** | 0.11** | 0.10** | 0.14** |
| non C>T | 0.13** | 0.11 | 0.13** | 0.19** |
| non C>A | 0.12** | 0.13 | 0.13** | 0.19** |
| non C>G | 0.12** | 0.17 | 0.31** | 0.24* |
| non T>C | 0.10** | 0.10 | 0.11** | 0.16* |
| non T>G | 0.0036 | 0.076** | 0.073** | 0.11 |
| non T>A | 0.090** | 0.082 | 0.081** | 0.12 |
| 100KB |  |  |  |  |
| All |  |  |  |  |
| CpG C>T | 0.068** | 0.044** | 0.043** | 0.061** |
| non C>T | 0.042** | 0.043 | 0.047** | 0.069** |
| non C>A | 0.049** | 0.046 | 0.043 | 0.072** |
| non C>G | 0.040** | 0.056 | 0.11** | 0.088** |
| non T>C | 0.041** | 0.041 | 0.034** | 0.063** |
| non T>G | 0.0064** | 0.028** | 0.023** | 0.044* |
| non T>A | 0.024** | 0.031 | 0.019** | 0.048** |

## Supplementary figures

**Figure S1.**
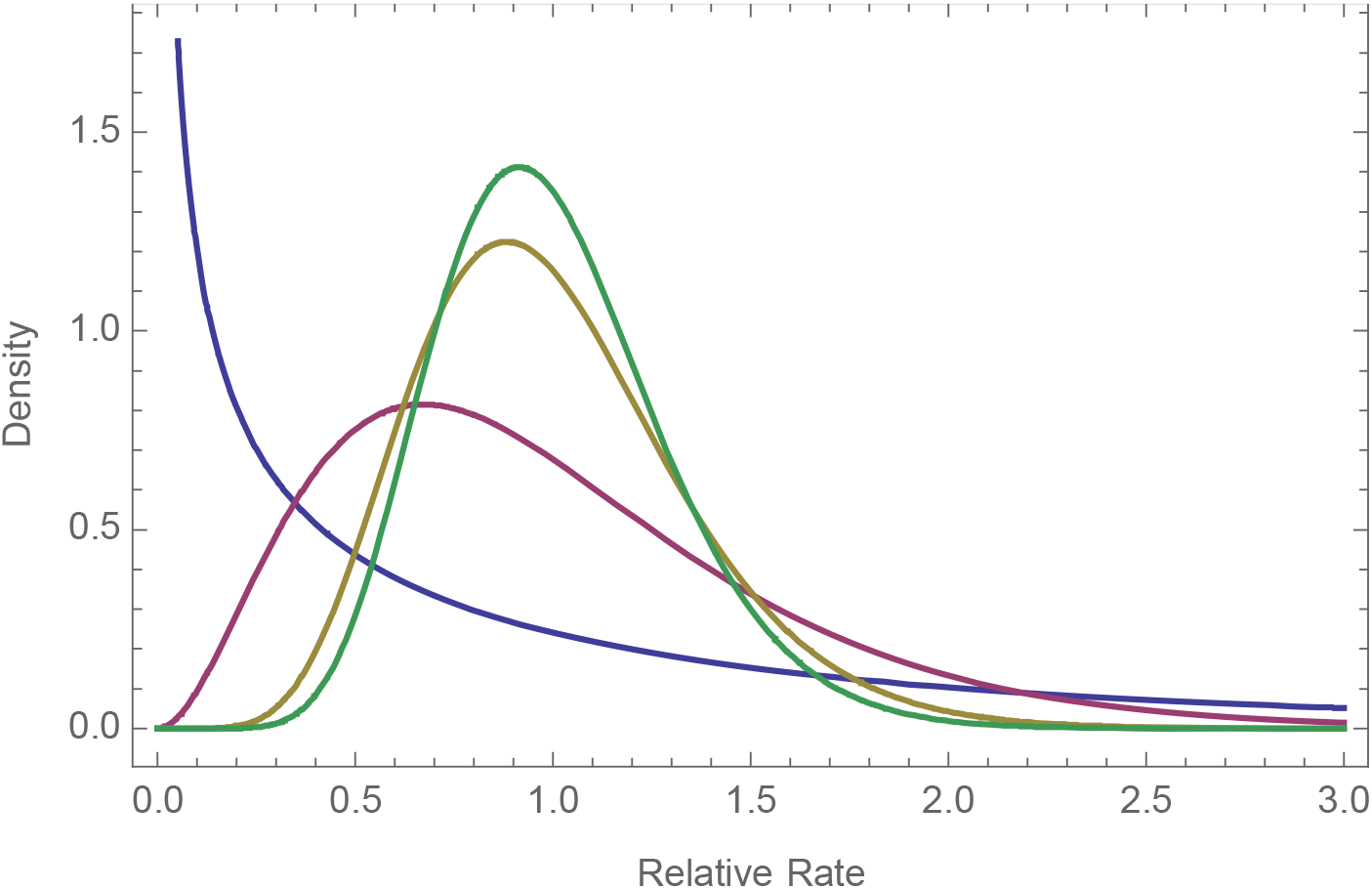
The gamma distribution fitted to the four datasets at the 100KB: Blue: Michaelson, Purple: Kong, Green: Francioli and Wong (coincident distributions), Olive: all data.

**Figure S2.**
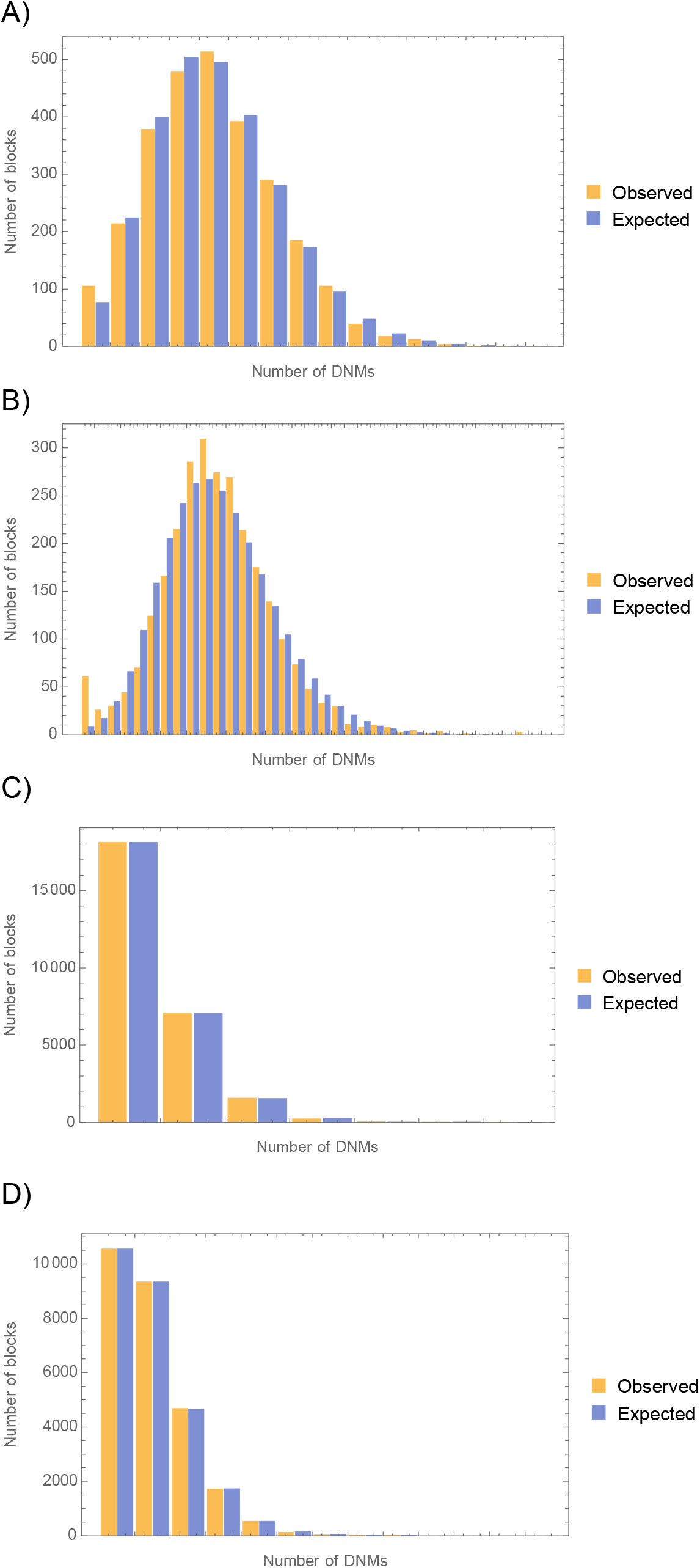
Goodness of fit of the gamma distribution. The distribution of observed and expected number of blocks with a given number of DNMs. The expected number was estimated using the fitted gamma distribution. A) Francioli 1MB, B) Wong 1MB, C) Francioli 100KB, and D) Wong 100KB. A goodness-of-fit test rejects the model for all datasets except Francioli 100KB.

**Figure S3.**
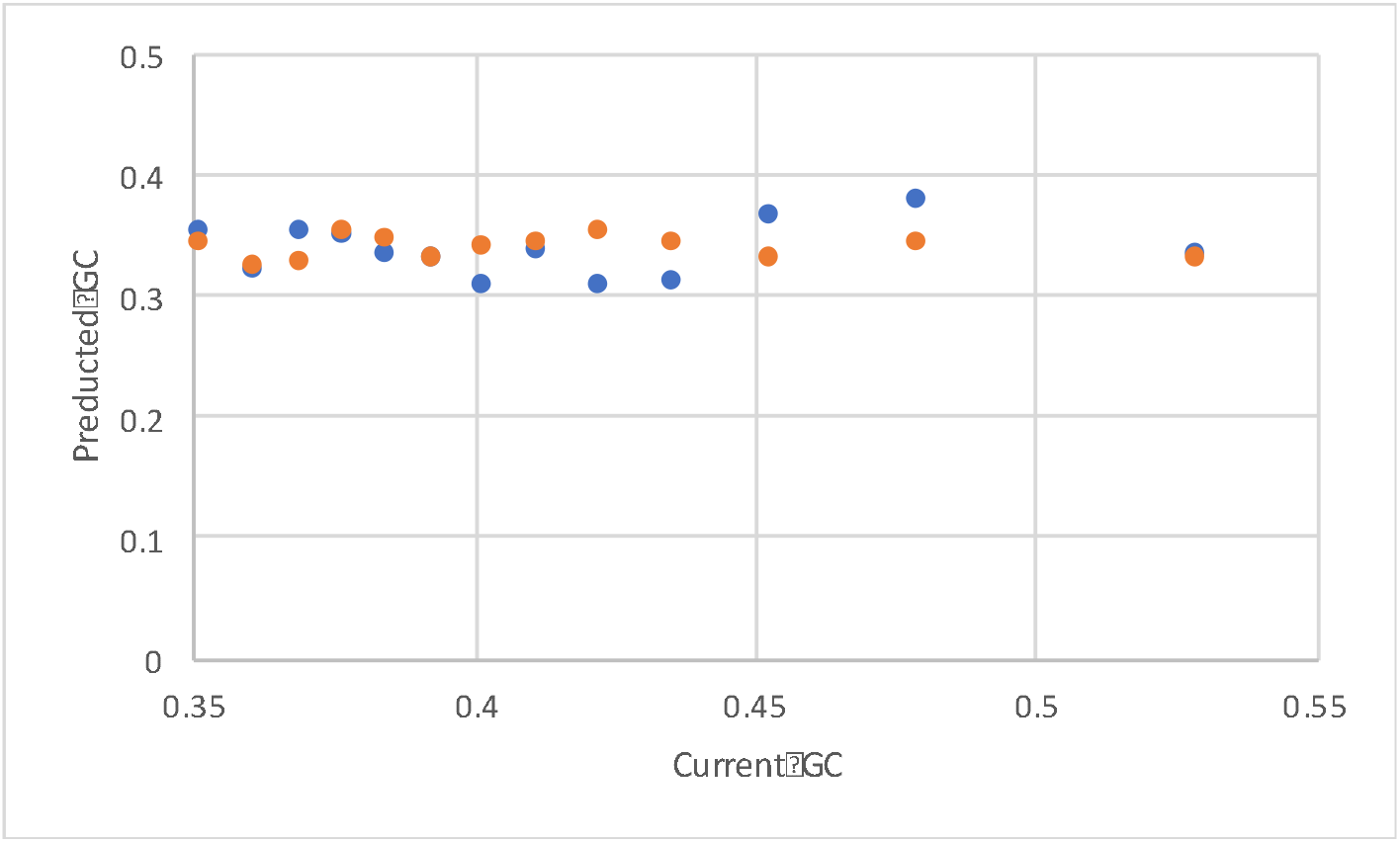
The predicted equilibrium GC content versus the current GC content using mutation rates inferred from the Francioli (blue) and Wong (orange) DNMs at the 100KB scale. The right-most data point is coincident.

**Figure S4.**
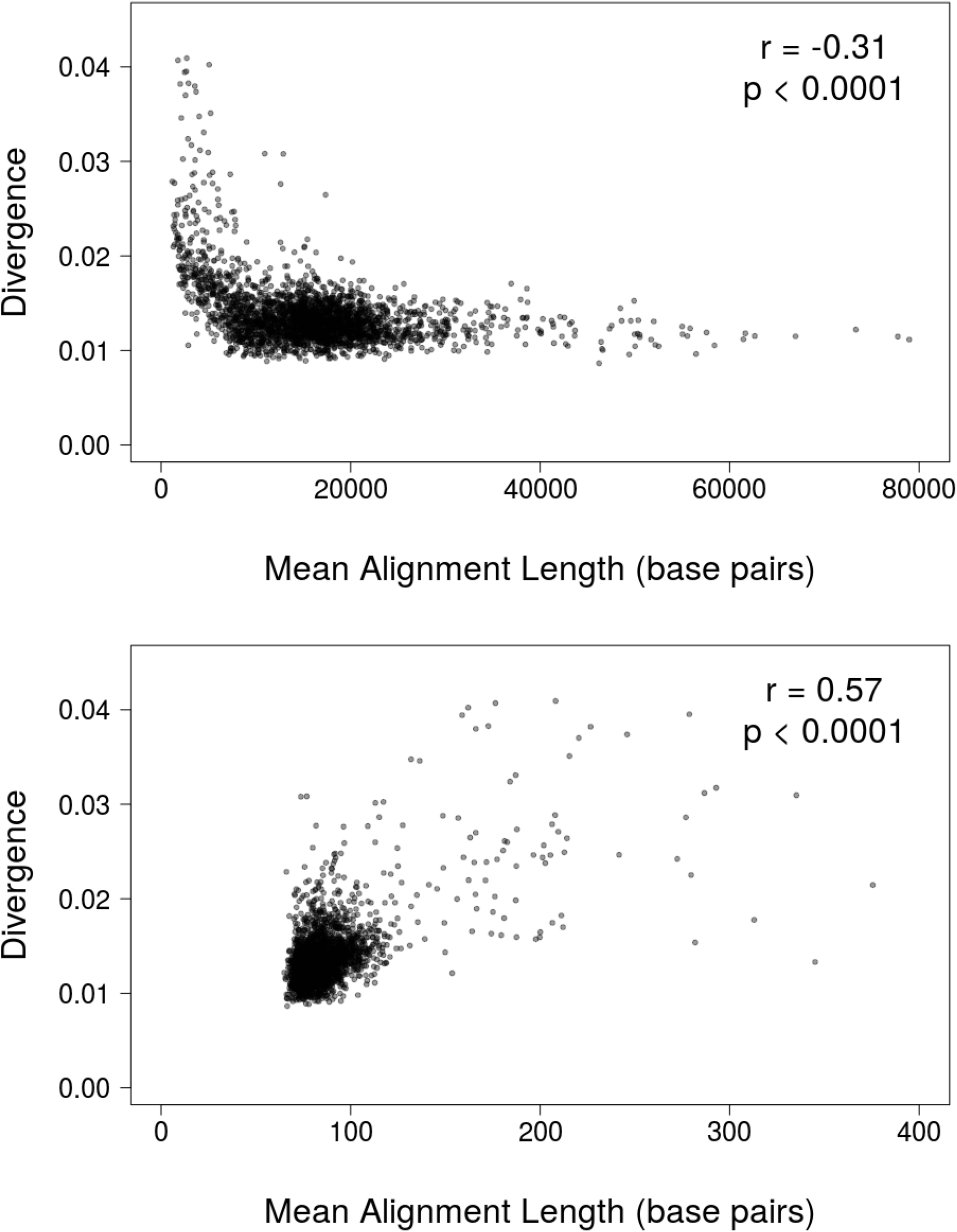
Divergence (number of substitutions per base pair) as a function of alignment length in the UCSD pairwise alignments (top panel) and the UCSD multiz alignments (bottom panel).

**Figure S5.**
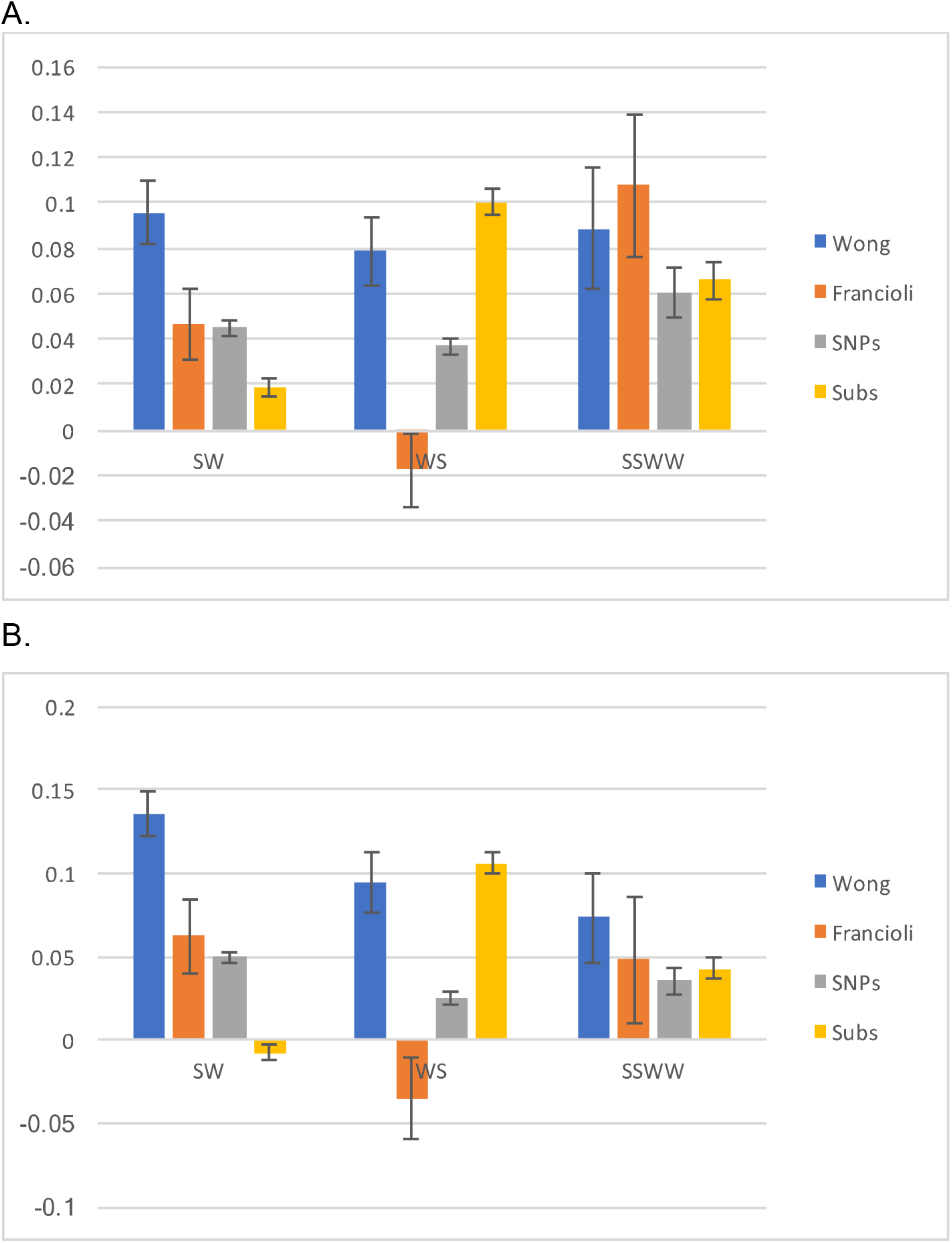
The slope (and SE) between normalised DNM density and normalised recombination rate (RR) (Wong - blue, Francioli - orange), normalised SNP density and RR (grey), and normalised substitution density and RR (yellow at 1MB scale. In each case the values were normalised by dividing the values by the mean. Panel A is for male recombination rates, panel B for female recombination rates.

**Figure S6.**
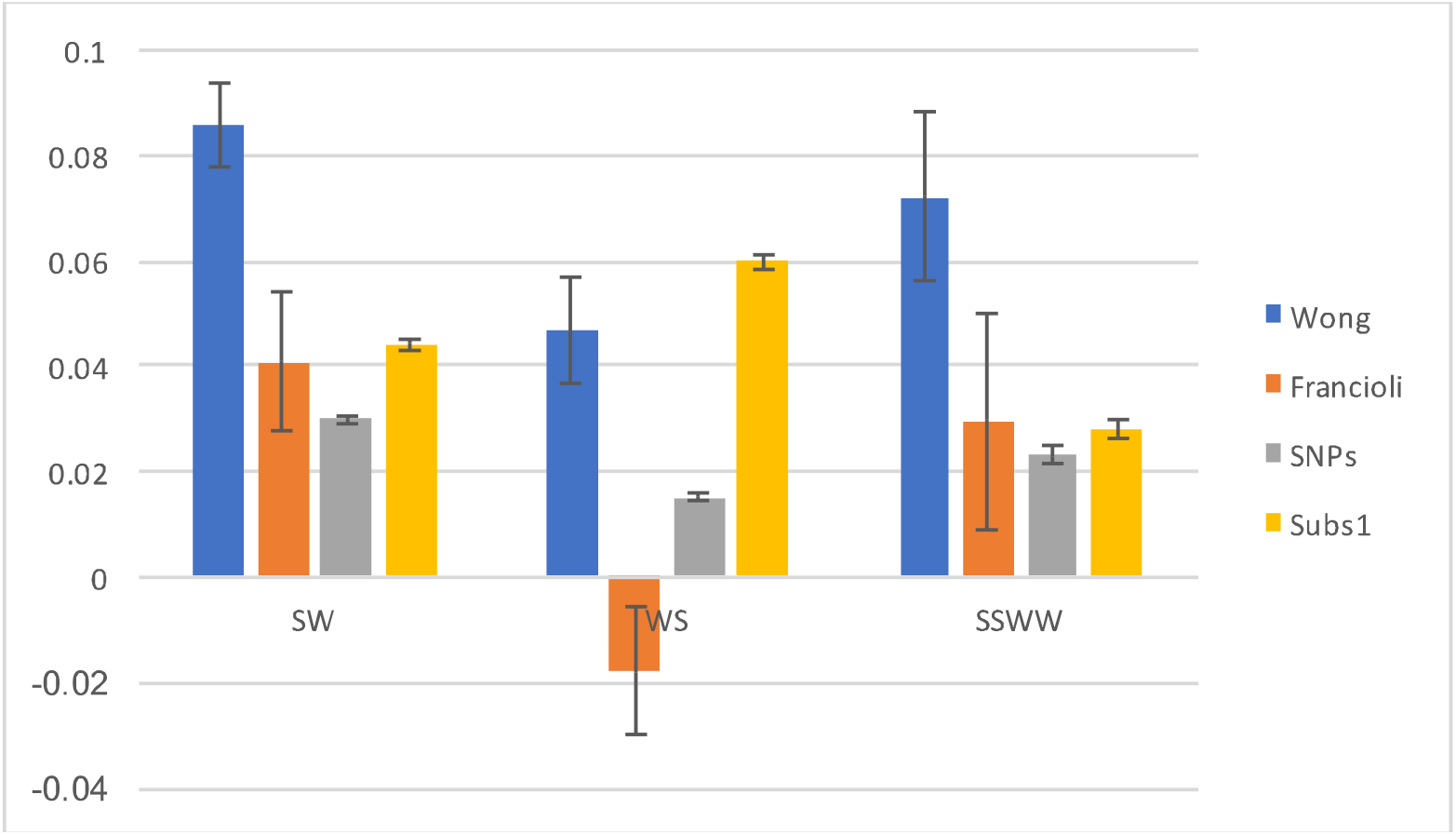
The slope between normalised DNM density and normalised recombination rate (RR) (Wong - blue, Francioli - orange), normalised SNP density and RR (grey), and normalised substitution density and RR (yellow at 100KB scale. In each case the values were normalised by dividing the values by the mean. Sex-averaged RRs were used.

**Figure S7.**
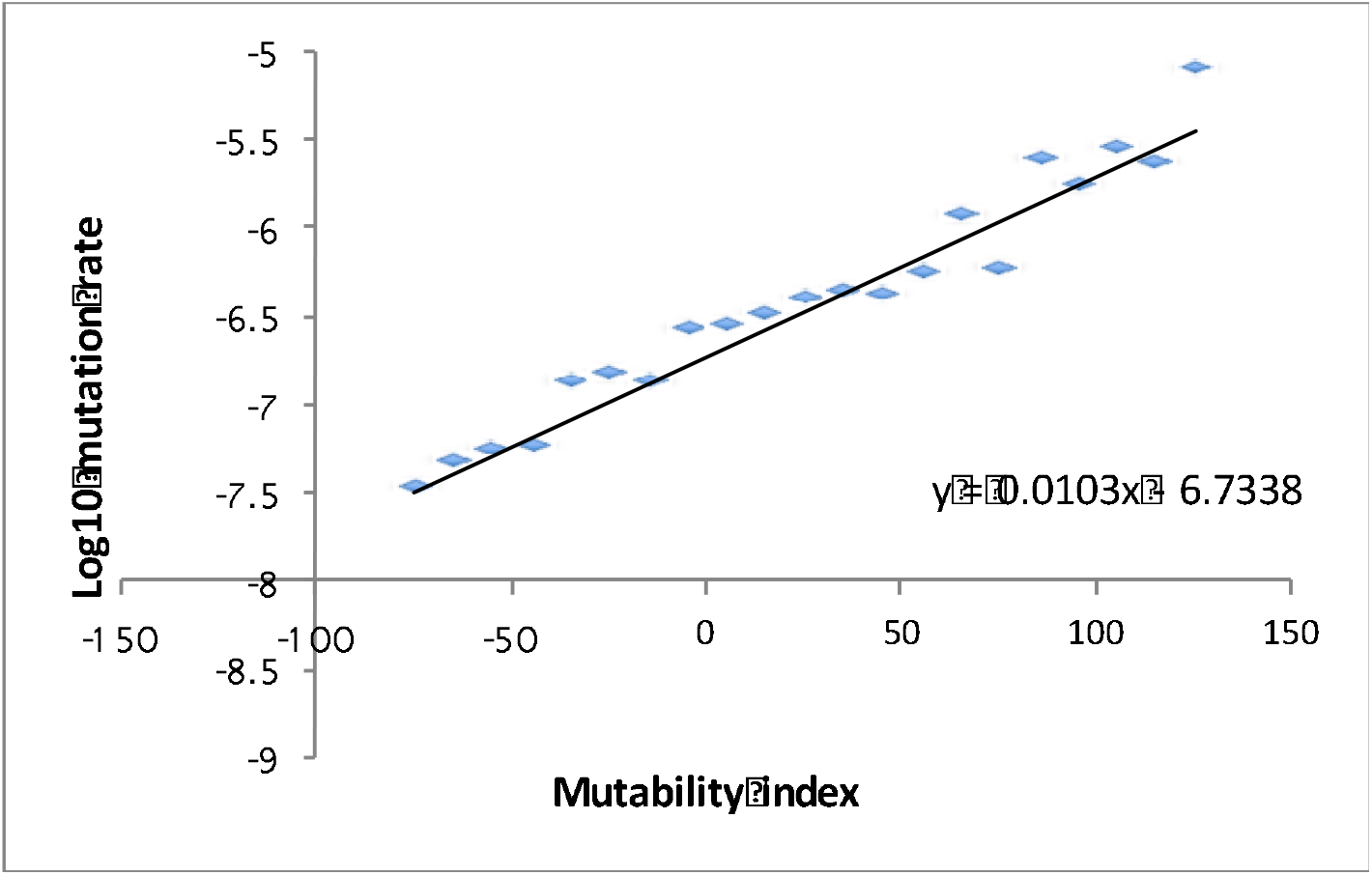
The relationship between the log mutation rate (estimated as number of DNMs over number of sites) and the mutability index from Michaelson et al.

